# Identifying potential key genes and existing drugs for Multiple sclerosis, Schizophrenia, and Autism- an in silico approach

**DOI:** 10.1101/2021.11.30.470019

**Authors:** Sajal Kumar Dey, Sushmita Bhowmick, Souvik Chakraborty

## Abstract

Nowadays, neurological conditions are a major concern as it not only preys on a patient’s health but also is a huge economic burden that is placed on the patient’s family. The diagnosis and treatment of disease sometimes cause methodological limitations. This is mainly common for individuals who have the signs of MS and schizophrenia (SZ). Patients suffering from multiple sclerosis are more likely to develop schizophrenia. Besides, a significant portion of patients who have been diagnosed with Autism Spectrum Disorder (ASD) later acquire the symptoms of Schizophrenia. In this study, we used bioinformatics tools to determine differentially expressed genes (DEGs) in all these diseases, and then we created a protein-protein interaction network using the online software STRING and identified 15 significant genes with the help of Cytohubba a plug-in tool in Cytoscape, the offline software (version3.8.2). We then used a drug-gene interaction database to conduct a drug-gene interaction study of the 15 hub genes and from there we identified 37 FDA-approved drugs. These findings may provide a new and common therapeutic approach for MS, SZ, and ASD therapy.

## 1. Introduction

Multiple sclerosis is an autoimmune disease in which cognitive problems of patients are commonly seen and the characteristic lesion in the central nervous system is also observed in the anatomical sections of specific areas of the brain of patients (Compston and Coles, 2008). It is estimated that 2.8 million people are believed to live with MS and females are more at risk than males (Walton et al., 2020). In MS, demyelination of the neurons occurs due to the activation of the innate branch of the immune system. Schizophrenia, an incurable disease characterized by psychotic delusional periods in patients which is triggered mainly by environmental and genetic factors. SZ can be seen as an interplay between the environmental and genetic factors which ultimately leads to the decrease in the cognitive ability of persons suffering from this dreadful disease (Kelly et al., 2021). Due to the poor understanding of the disease, symptomatic treatments are right now the best we got for the patients (Stępnicki et al., 2018). Due to their etiological similarities, it is thought that both MS and SZ might be caused by the same mechanisms but MS and SZ are different with respect to the fact that MS is caused by immune responses in the specific areas of the brain but SZ is altogether a different ball game that starts as hallucinogenic episodes and ultimately leads to psychotic changes in the behavior of patients. (Arneth, 2017). Autism spectrum disorders (ASDs) is an umbrella term given to a series of behavioral disorders related to social interaction, reduced mental abilities, and also restricted patterns of interests. It is essentially a developmental disorder of the central nervous system (Memari et al., 2015). The cases for ASDs are steadily increasing and are predominant in children less than 5 years of age (Chiarotti and Venerosi, 2020). Once thought to be a predecessor of SZ, ASDs and SZ are now considered two very much different diseases though some studies now show that individuals with ASDs are more inclined towards SZ (De Crescenzo et al., 2019). Genetic contribution in the case of SZ is about 25–33% and for ASD this goes up to 49%, which shows that both SZ and ASDs have elements of genetic interplay for the development of the disease (St Pourcain et al., 2018).

In this study, we applied bioinformatics tools to determine the common genes and pathways associated with MS, SZ, and ASD based on publicly accessible microarray data from gene expression omnibus (Edgar et al., 2002). Then we uploaded those genes for protein-protein interaction analysis in the software String and imported the String network into the software Cytoscape to determine the 15 hub genes. Furthermore, we conducted a drug-gene interaction study of hub genes using the drug-gene interaction database (DGIdb), which may help in matching certain available drugs and, ultimately, discovering alternatives for prevention and therapy.

## 2. Material and methods

### 2.1 Retrieval of microarray data

All the datasets were retrieved from the gene expression omnibus (GEO) database of NCBI (https://www.ncbi.nlm.nih.gov/geo/) by using the keywords multiple sclerosis, schizophrenia, and autism respectively. After careful consideration, three gene expression profiles (GSE21942, GSE17612, GSE25507) were downloaded from the GEO datasets database. The microarray gene expression data for MS is GSE21942 which was based on platform GPL570 [HG-U133_Plus_2] Affymetrix Human Genome U133 Plus 2.0 Array. GSE17612, the microarray gene expression data for SZ was based on platform GPL570 [HG-U133_Plus_2] Affymetrix Human Genome U133 Plus 2.0 Array. GSE25507, the gene expression data for ASD was based on platform GPL570 [HG-U133_Plus_2] Affymetrix Human Genome U133 Plus 2.0 Array. The gene expression profile GSE21942 (MS) contains a total of 27 samples and from that 12 test samples and 15 control samples were selected. GSE17612 (SZ) contains a total of 51 samples. From that total, 28 control samples and 23 test samples were selected. In GSE25507, a total of 146 samples are present. From that total, 84 samples were defined as tests, and 64 samples were selected as control.

### 2.2 Data processing and identification of DEGs

Significant DEGs between test and control samples were analyzed by using an online analysis tool GEO2R (https://www.ncbi.nlm.nih.gov/geo/geo2r/) for each dataset. The GEO2R tool uses Bioconductor package (GEOQuery and limma) for the processing of those data(Clough and Barrett, 2016). DEGs were screened with the following cut-off criteria: P-value < 0.05 and a fold change value (logFC) greater than equal to 0.5 for each dataset.

### 2.3 Identification of common DEGs

After screening, all the DEGs for each dataset were uploaded in a desktop-based software Funrich (http://www.funrich.org/) (Pathan et al., 2015). By using this Funrich software, intersecting DEGs were obtained via the Venn diagram.

### 2.4 Network analysis of common DEGs

After getting common DEGs from Funrich software, they were imported into an online-based software GeneMANIA (https://genemania.org/) for generating a hypothesis about gene function and for the development of a gene network(Franz et al., 2018).

All the DEGs were uploaded into the online-based tool String (https://string-db.org/) for protein-protein interaction (PPI) analysis. The String database collects all protein-protein interaction information from all publicly available sources and its goal is to create a network that includes both physical and functional interactions (szklarczyk et al., 2019_). The PPI network was generated with an interaction score of 0.4 and all disconnected nodes in the network were hidden.

### 2.5 Identification of hub genes

The PPI network retrieved from the online software String was then imported into a desktopbased software Cytoscape (https://cytoscape.org/). The top 15 genes were identified via using a plugin tool Cytohubba in the software Cytoscape by using the degree method. These top 15 genes in the PPI network were termed hub genes

### 2.6 Analysis of GO and KEGG pathway of DEGs

For further study, we uploaded all the DEGs into the web-based tool Enrichr (https://maayanlab.cloud/Enrichr/) for the enrichment analysis. In the software Enrichr, gene functions and the KEGG pathway were obtained. Gene function can be classified into biological process, molecular function, and cellular component. All three functions of the gene were obtained via the software.

### 2.7 microRNA network analysis

The top 15 hub genes were submitted to the online software miRNet (https://www.mirnet.ca/) to find out the interaction between miRNAs and genes. A network between genes and miRNAs was generated by using the software.

### 2.8 Analysis of Drug-gene interaction

To detect the relationship between hub genes and existing drugs we imported the hub genes into the drug-gene interaction database (DGIdb https://www.dgidb.org/) to search for existing agonist, antagonist, and inhibitor drugs that are FDA approved. After downloading the file from DGIdb, we screened the data. After screening, the data was imported into a computer-based software Cytoscape (version3.8.2) (https://cytoscape.org/) for visualization. After selecting nodes and edges, the interaction between hub genes and drugs was visualized in the above-mentioned software Cytoscape.

## 3. Results

### 3.1 Identification of DEGs

In this study, three gene expression profiles (GSE21942, GSE17612, GSE25507) were selected. The differential expression analysis showed the dataset GSE21942 contains 1305 upregulated genes that were obtained based on the following cut-off criteria: P-value < 0.05 and a fold change value (logFC) greater than equal to 0.5, as well as 246 and 1593 upregulated genes were identified in the datasets GSE17612 and GSE25507 respectively.

### 3.2 Identification of common DEGs

All the upregulated genes from each dataset were uploaded to the desktop-based software Funrich. To find the intersection of the DEGs, a Venn analysis was performed. Two genes are common among the three datasets. 155 genes are common between the datasets GSE21942 (MS) and GSE25507 (ASD), 25 and 12 upregulated genes are common between the datasets GSE17612 (SZ) and GSE25507 (ASD), GSE21942 (MS), and GSE17612 (SZ) respectively. The following diagram is the demonstration of the common DEGs among the three diseases (Fig 1).

**Fig 1:**
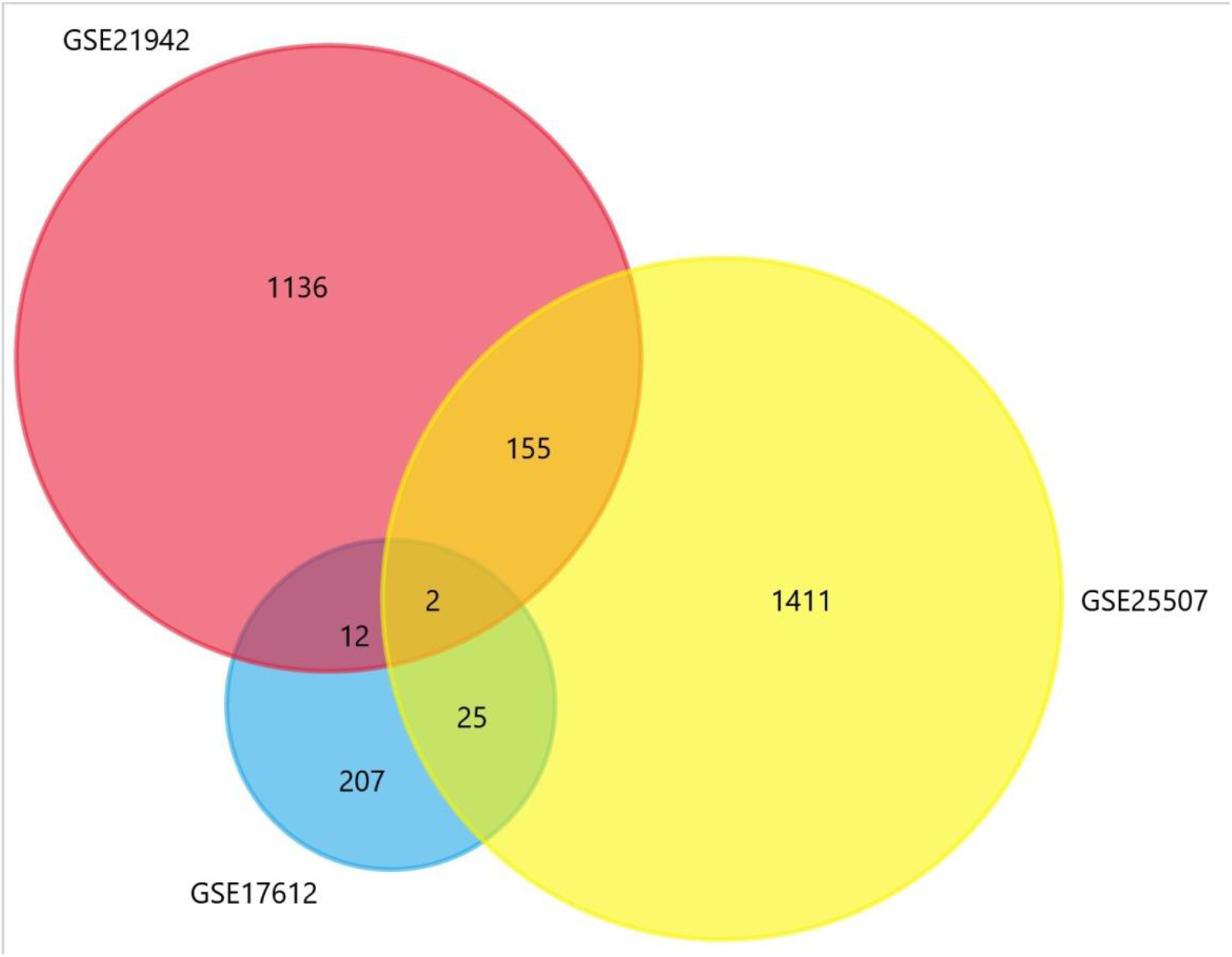
Venn diagram represents the intersecting upregulated DEGs among the three datasets GSE21942 (MS), GSE17612 (SZ), and GSE25507 (ASD).

### 3.3 Network analysis of common DEGs

#### 3.3.1 GeneMANIA

GeneMANIA, an online software was used to predict the interactions among the common DEGs. Here, the circles represent the genes, and the lines represent the interactions among the DEGs (Fig 2). The pink lines represent the physical interaction among them, the green and violet lines represent genetic interaction and coexpression respectively.

**Fig 2:**
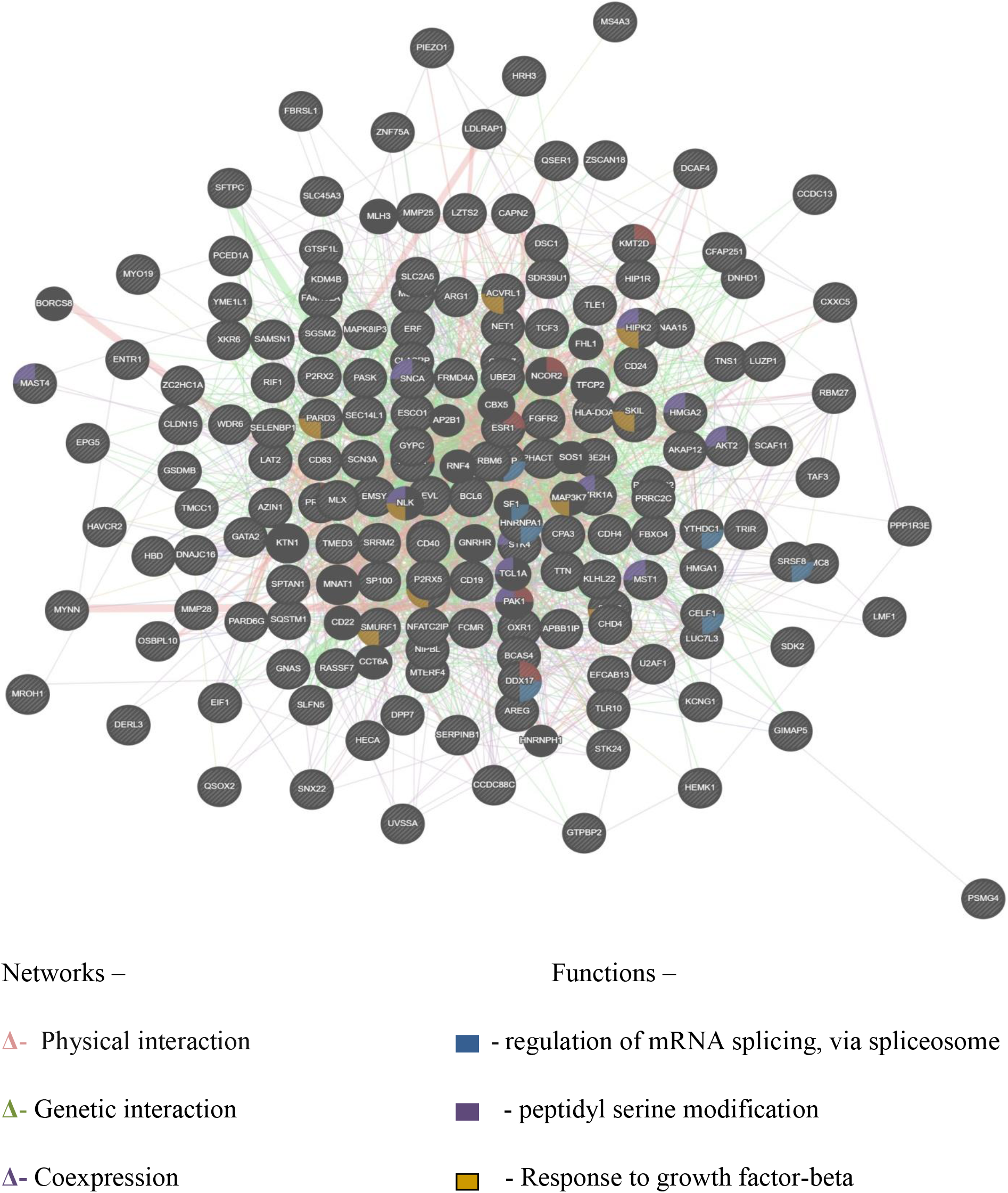
Network between common DEGs, created using GeneMANIA software

#### 3.3.2 Protein-protein interaction analysis

With the help of STRING, an online software protein interactions among the DEGs were predicted. A total no of 168 nodes and 147 edges were involved in the predicted PPI network (Fig 3).

**Fig 3:**
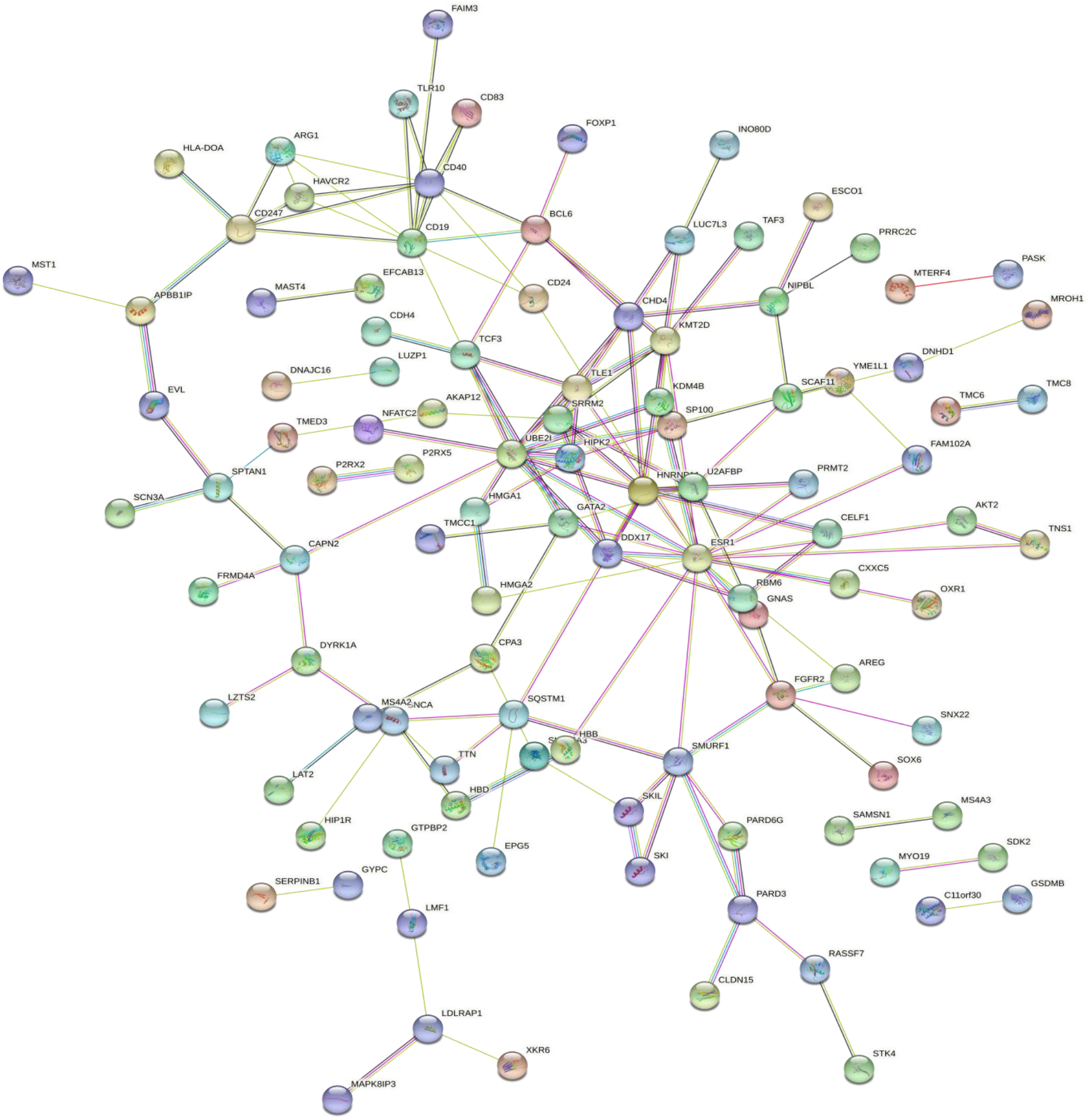
STRING protein-protein interaction network of 194 upregulated genes. Here circles represent the genes and lines represent the interaction between them.

#### 3.4 Analysis of hub genes

We used Cytohubba, a plug-in tool in the Cytoscape to identify the top 15 genes from the PPI network which we got from the software String(Fig 4). It had been seen that the gene, Estrogen receptor 1 (ESR1) possesses the highest degree with a score of 19. UBE2I, CD19, HNRNPA1, CD40, and SMURF1 have degree scores of 12, 10, 9, 8, and 7 respectively. The genes SRRM2, CD247, FGFR2, BCL6, TCF3, KMT2D, and CHD4 possess a degree score of 6. Gata binding factor 2 (GATA2) and dead box helicase 17 (DDX17) have the lowest degree with a score of 5. In the following diagram, the color code represents the degree. The red, orange and yellow colors denote the highest, medium, and lowest degree of centrality(Fig 4).

**Fig 4:**
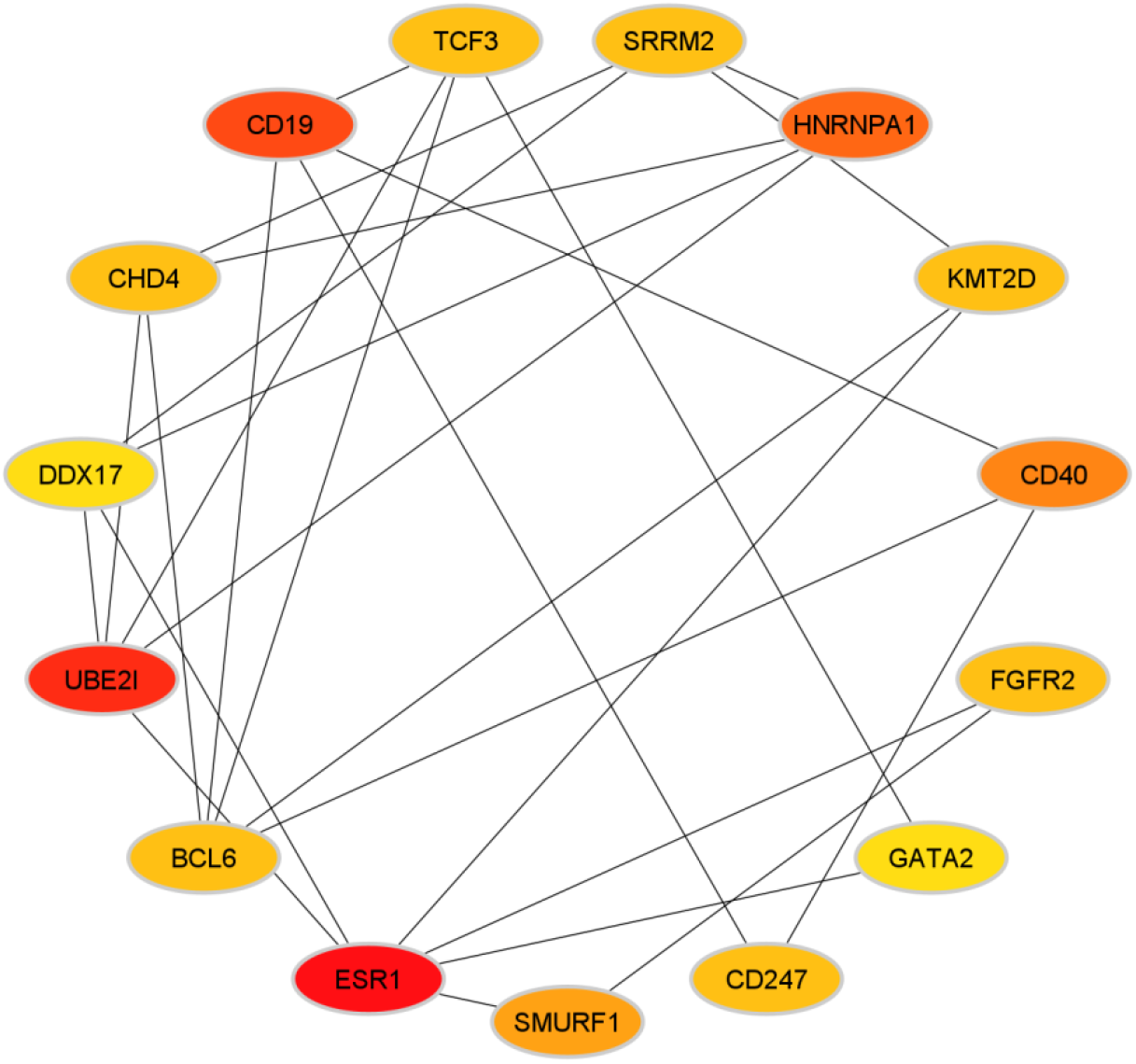
Presentation of hub genes using a plug-in tool Cytohubba in the offline software Cytoscape. Here the circles represent the name of the genes and the lines represent the interactions between them. We generated this diagram using the degree method. Here the color represents the degree.

#### 3.5 Functional enrichment analysis of DEGs

By using the online software Enrichr, GO functions and KEGG pathway enrichment analysis were performed for upregulated DEGs. After submitting, all the DEGs in the online software Enrichr, under the ontology section several options were given. Among them GO biological process (BP), GO molecular function(MF) and GO cellular component(CC) was identified. The BP analysis showed that the DEGs were enriched in negative regulation of transcription by RNA polymerase II, positive regulation of carbohydrate metabolic process and calcium iron transport into the cytosol and also the small molecule metabolic process, regulation of BMP signaling pathway, spliceosome complex assembly, histone-serine phosphorylation and regulation of mRNA splicing (Fig 5a). For MF analysis, the DEGs were enriched in C2H2 zinc finger domain binding, hemoglobin alpha binding, chromo shadow domain binding, the minor groove of adenine-thymine rich DNA binding, nucleotide receptor activity, peroxisome proliferator-activated receptor binding, extracellularly ATP gated cation channel activity, protein serine/threonine kinase activity and pre mRNA binding (Fig 5b). The CC analysis showed that the DEGs were enriched in integral and intrinsic components of the nuclear inner membrane, specific granule, bicellular tight junction, NatA complex, axon, clathrin-coated endocytic vesicles, alpha-beta T cell receptor complex, and alveolar lamellar body (Fig 5c). The analysis of the KEGG pathway showed that the DEGs are mainly enriched in asthma, rap-1 signaling pathway, malaria, signaling pathway regulating pluripotency of stem cells, insulin resistance, toxoplasmosis, Fc epsilon R1 signaling pathway, non-small cell lung cancer, FoxO signaling pathway, and transcriptional misregulation in cancer.

**Fig 5:**
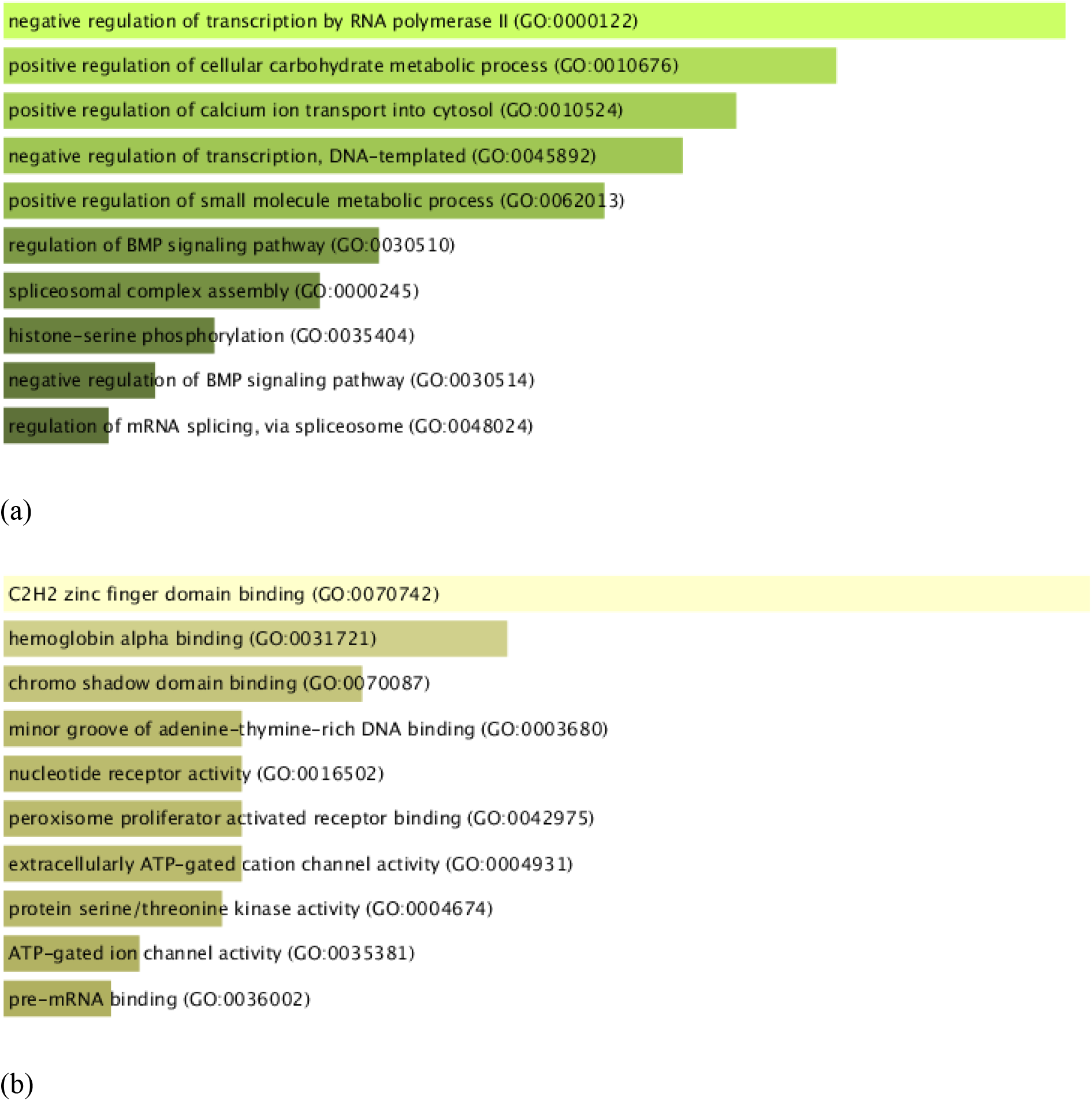
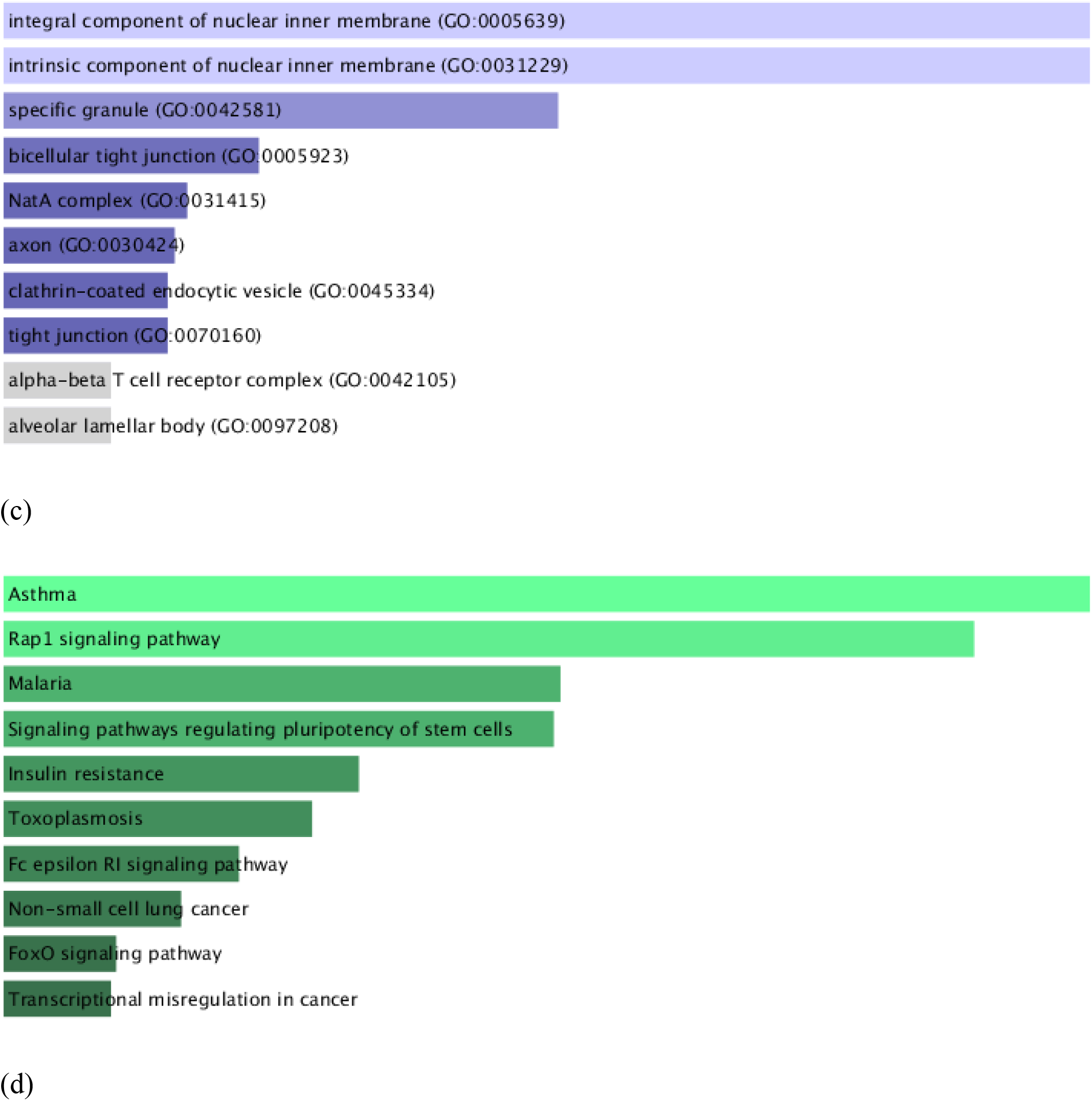
GO function and KEGG pathway analysis by using the online software Enrichr. (a)A top ten enriched biological processes for DEGs. The X and Y-axis represent the gene and the biological process respectively. (b) The top ten enriched molecular functions for DEGs. The X-axis represents the gene name and the Y-axis represents the molecular function (c) The top ten enriched cellular components for DEGs. The X-axis represents the gene name and the Y-axis represents the cellular components. (d) The top ten KEGG pathways for DEGs. The X and Y-axis represent the gene and the KEGG pathway respectively.

**Fig 6:**
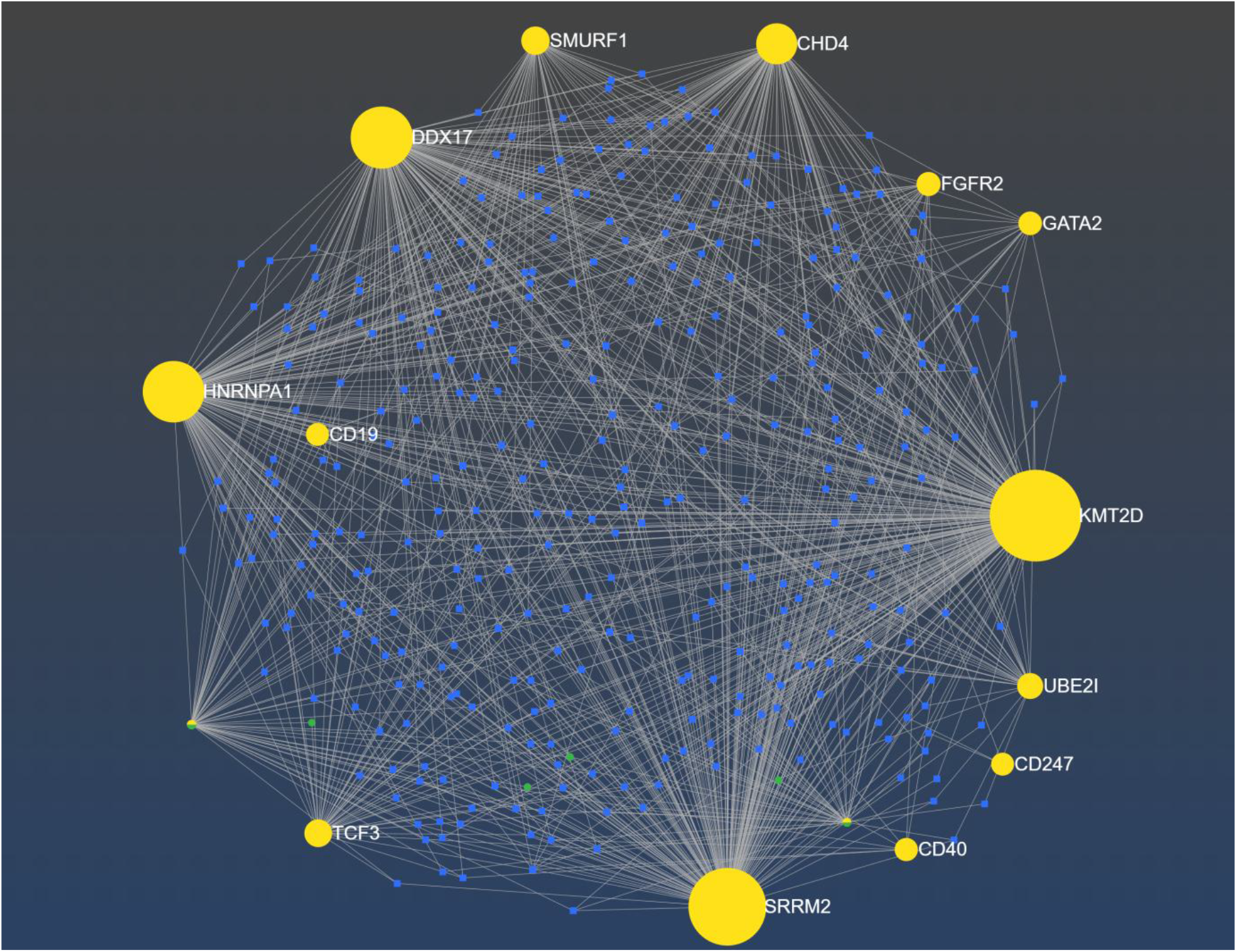
Interaction among DEGs and miRNAs were generated by using the online software miRNet. The genes, miRNAs, and TF are represented in yellow, blue, and green colors respectively.

#### 3.6 microRNA interaction analysis

After uploading all 15 hub genes in the online software, a network between gene and miRNAs were generated. Six transcription factors (TFs) and 339 miRNAs were found in association with 15 hub genes. In the network, the gene KMT2D with maximal connectivity possesses the highest degree score of 205, followed by the gene SRRM2 with a 181 degree score. The genes DDX17, HNRNPA1, CHD4, ESR1, SMURF1, TCF3, BCL6, UBE2I, FGFR2, and GATA2 possess the degree score of 148, 137, 98, 58, 51, 50, 40, 37, 21, and 18.

#### 3.7 Analysis of Drug-gene interaction

After uploading 15 hub genes in the drug-gene interaction database we got FDA-approved drugs against those genes. We downloaded the data as a tsv file then we selected only the agonist, antagonist, and inhibitor drugs and got 37 FDA-approved drugs. We got 28 drugs for ESR1 and 9 drugs for FGFR2. For visualization, we uploaded the screened data into the software Cytoscape and generated two images to present the agonist, antagonist, and inhibitors against the genes ESR1 (Fig 7a) and FGFR2 (Fig 7b) respectively.

**Fig 7:**
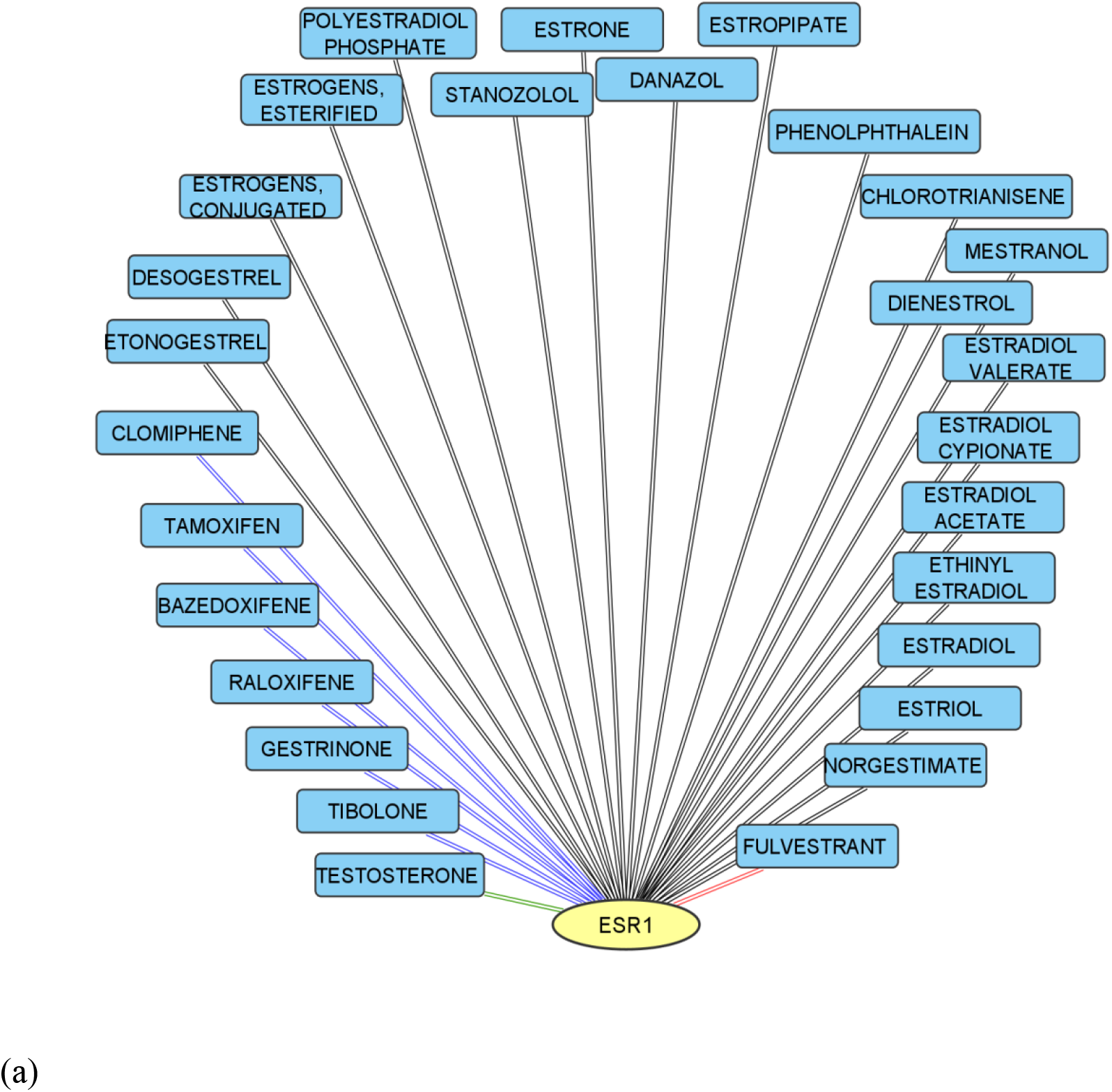
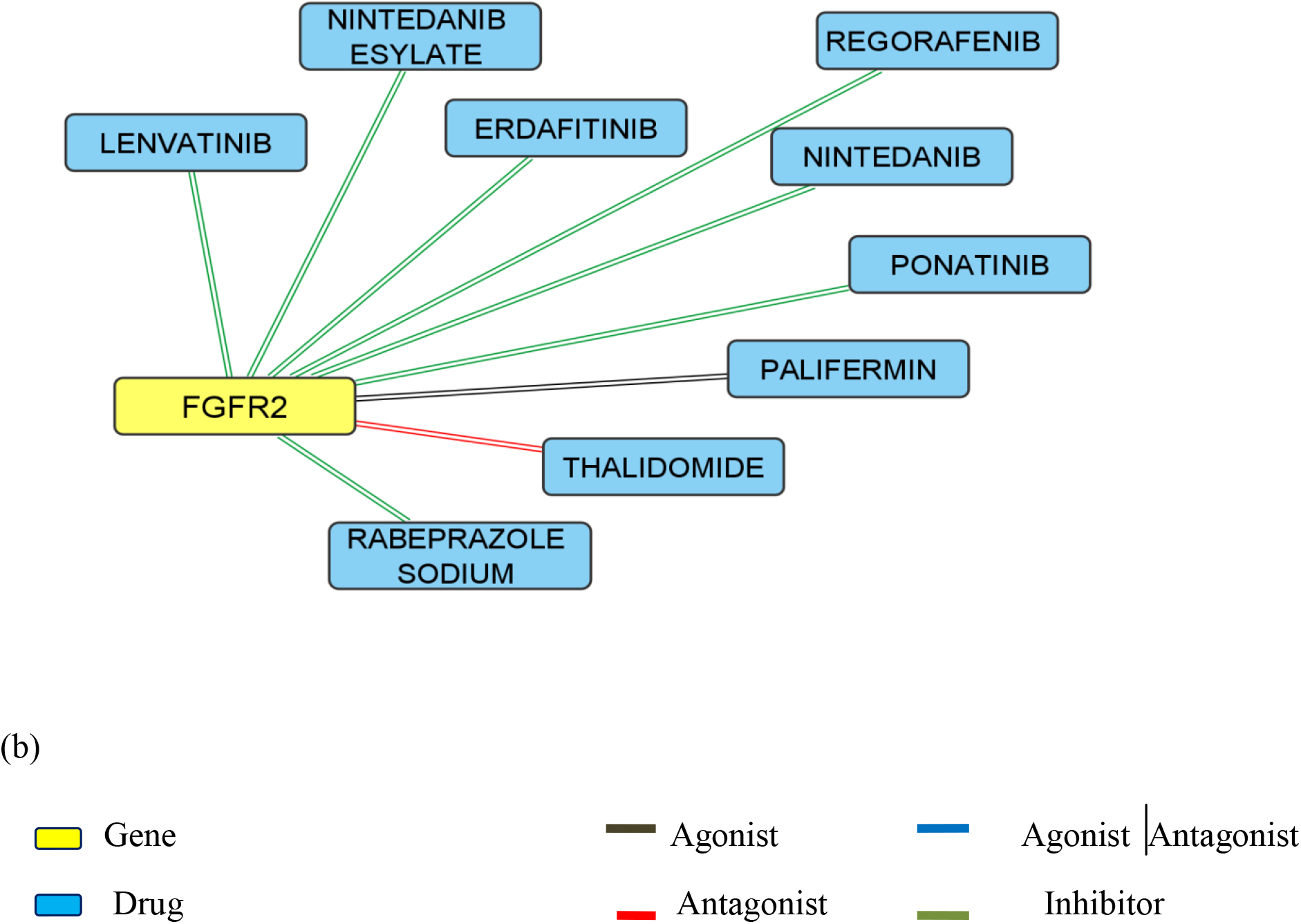
Visualisation of Drug-gene interaction using the offline software Cytoscape (a) drugs for gene ESR1 (b) drugs for gene FGFR2.

## 4. Discussion

The pathway and genes which are involved in neurological disorders are very complex. In this analysis, we utilized three datasets for three different neurological disorders (MS, SZ, and ASD) to look for the similarities in the expression of genes.

### 4.1 Analysis of network

After finding the common genes among these three datasets by using the software Funrich we generated a PPI network and then to find the top 15 genes we imported the PPI network to the desktop-based software Cytoscape. Those top 15 genes are ESR1, UBE2I, CD19, HNRNPA1, CD40, SMURF1, SRRM2, CD247, FGFR2, BCL6, TCF3, KMT2D, CHD4, GATA2 andDDX17. Among these hub genes ESR1, UBE2I and cd19 possess a higher degree score.

The gene ESR1 which possesses the highest degree score encodes the protein named estrogen receptor, a nuclear hormone receptor activated by the hormone estrogen. There are two estrogen receptor subtypes present in the human forebrain α and β. The ERβ is present in the hippocampus, thalamus, and entorhinal cortex, suggesting it plays an important role in cognition, mood, memory formation, and motor functions(Österlund and Hurd, 2001). The expression of ERα mRNA appears to play a significant role in the hypothalamus and amygdala which is associated with emotion(Österlund and Hurd, 2001). Besides these, estrogen helps in the regulation of neurotransmissions such as serotonin, norepinephrine, and acetylcholine(Sundermann et al., 2010). The sites which are associated with the estrogen receptor are highly associated with these three neurological conditions (MS, SZ, and ASD).

Ubiquitin-conjugating enzyme E2I (UBE2I) is a protein-coding gene that encodes SUMO-conjugating enzyme UBC9 which accepts the small ubiquitin-related modifier (SUMO) and with the help of E3 ligase, it helps in the attachment of ubiquitin-like proteins to the target proteins, resulting in the changes of the target protein’s activity and localization(Ahn et al., 2009).

The gene CD19 encodes the protein, B lymphocyte antigen CD19, a type I transmembrane glycoprotein is expressed throughout the B cell development(Stüve et al., 2014). Upon the activation of B cell receptors, CD19 through the activation of PI3 and subsequently Akt kinases, enhances BCR induced signaling crucial for the expansion of B cells(Chung et al., 2012).

### 4.2 Drug-gene interaction analysis

After finding the top 15 genes, we imported those genes into the drug-gene interaction database and 37 existing drugs were found. Among those 37 drugs, there there are 28 drugs for ESR1 and 9 drugs for FGFR2. Among those 28 drugs for ESR1, Fulvestrant acts as an antagonist and the drug Tibolone, Gestrinone, Raloxifene, Bazdoxifene, Tamoxifen, and Clomiphene have both agonists as well as antagonists characteristics.

3-α and 3-β-OH Tibolone act as an antagonist for ESR1(Erβ) but interestingly Δ4 Tibolone and Tibolone possess agonist activity for ESR1(Escande et al., 2009). Tamoxifen act as a neuroprotective drug like the sex hormone estrogen. Estrogen reduces neuronal cell death by blocking oxidative stress(Ernst et al., 2002). Clomiphene possesses agonist activity for ESR1 but in higher doses, the antagonistic activity of Clomiphene has also been shown(Bowman et al., 1981). Raloxifene and Bazdoxifene both are selective estrogen modulators. They both have agonist activity in certain tissue and antagonist activity in certain tissue. Due to their tissue selectivity, they both have fewer side effects(Johnson and Hauck, 2016) (Scott et al., 1999).

Among the 9 drugs for FGFR2, Thalidomide acts as an antagonist and Palifermin acts as an agonist and the other drugs are Lenvatinib, Rabeprazole sodium, Ponatinib, Nintedanib, Regorafenib, Erdafitinib, and Nintedinib esylate act as an inhibitor.

## 5. Conclusion

In conclusion, we identified some key genes that are associated with these three neurological disorders (MS, SZ, and ASD). Some genes had not been previously reported but might play an important role in these three neurological disorders. Among these key genes, ESR1, UBE2I, and CD19 possess higher degree scores and we found the existing drugs against genes ESR1 and FGFR2.

## Supporting information

Gene mania report

STRING DATABASE

Micro RNA enrichment

## References

Ahn, K., Song, J.H., Kim, D.K., Park, M.H., Jo, S.A., Koh, Y.H., 2009. Ubc9 gene polymorphisms and late-onset Alzheimer’s disease in the Korean population: A genetic association study. Neuroscience Letters 465, 272–275. https://doi.org/10.1016/j.neulet.2009.09.017

Arneth, B.M., 2017. Multiple Sclerosis and Schizophrenia. Int J Mol Sci 18, 1760. https://doi.org/10.3390/ijms18081760

Bowman, S.P., Leake, A., Miller, M., Morris, I.D., 1981. AGONIST AND ANTAGONIST ACTIVITY OF EN-CLOMIPHENE UPON OESTROGEN-MEDIATED EVENTS IN THE UTERUS, PITUITARY GLAND AND BRAIN OF THE RAT. Journal of Endocrinology 88, 367–374. https://doi.org/10.1677/joe.0.0880367

Chiarotti, F., Venerosi, A., 2020. Epidemiology of Autism Spectrum Disorders: A Review of Worldwide Prevalence Estimates Since 2014. Brain Sciences 10, 274. https://doi.org/10.3390/brainsci10050274

Chung, E.Y., Psathas, J.N., Yu, D., Li, Y., Weiss, M.J., Thomas-Tikhonenko, A., 2012. CD19 is a major B cell receptor–independent activator of MYC-driven B-lymphomagenesis. J Clin Invest 122, 2257–2266. https://doi.org/10.1172/JCI45851

Clough, E., Barrett, T., 2016. The Gene Expression Omnibus database. Methods Mol Biol 1418, 93–110. https://doi.org/10.1007/978-1-4939-3578-9_5

Compston, A., Coles, A., 2008. Multiple sclerosis. The Lancet 372, 1502–1517. https://doi.org/10.1016/S0140-6736(08)61620-7

De Crescenzo, F., Postorino, V., Siracusano, M., Riccioni, A., Armando, M., Curatolo, P., Mazzone, L., 2019. Autistic Symptoms in Schizophrenia Spectrum Disorders: A Systematic Review and Meta-Analysis. Front Psychiatry 10, 78. https://doi.org/10.3389/fpsyt.2019.00078

Edgar, R., Domrachev, M., Lash, A.E., 2002. Gene Expression Omnibus: NCBI gene expression and hybridization array data repository. Nucleic Acids Research 30, 207–210. https://doi.org/10.1093/nar/30.L207

Ernst, T., Chang, L., Cooray, D., Salvador, C., Jovicich, J., Walot, I., Boone, K., Chlebowski, R., 2002. The Effects of Tamoxifen and Estrogen on Brain Metabolism in Elderly Women. JNCI: Journal of the National Cancer Institute 94, 592–597. https://doi.org/10.1093/jnci/94.8.592

Escande, A., Servant, N., Rabenoelina, F., Auzou, G., Kloosterboer, H., Cavaillès, V., Balaguer, P., Maudelonde, T., 2009. Regulation of activities of steroid hormone receptors by tibolone and its primary metabolites. The Journal of Steroid Biochemistry and Molecular Biology 116, 8–14. https://doi.org/10.1016/j.jsbmb.2009.03.008

Franz, M., Rodriguez, H., Lopes, C., Zuberi, K., Montojo, J., Bader, G.D., Morris, Q., 2018. GeneMANIA update 2018. Nucleic Acids Res 46, W60–W64. https://doi.org/10.1093/nar/gky311

FunRich: An open access standalone functional enrichment and interaction network analysis tool - Pathan - 2015 - PROTEOMICS - Wiley Online Library [WWW Document], n.d. URL https://analyticalsciencejournals.onlinelibrary.wiley.com/doi/10.1002/pmic.201400515 (accessed 9.28.21).

Johnson, K., Hauck, F., 2016. Conjugated Estrogens/Bazedoxifene (Duavee) for Menopausal Symptoms. AFP 93, 307–314.

Kelly, J.R., Minuto, C., Cryan, J.F., Clarke, G., Dinan, T.G., 2021. The role of the gut microbiome in the development of schizophrenia. Schizophrenia Research, The Microbiome in Schizophrenia 234, 4–23. https://doi.org/10.1016/j.schres.2020.02.010

Memari, A.H., Panahi, N., Ranjbar, E., Moshayedi, P., Shafiei, M., Kordi, R., Ziaee, V., 2015. Children with Autism Spectrum Disorder and Patterns of Participation in Daily Physical and Play Activities. Neurology Research International 2015, e531906. https://doi.org/10.1155/2015/531906

Österlund, M.K., Hurd, Y.L., 2001. Estrogen receptors in the human forebrain and the relation to neuropsychiatric disorders. Progress in Neurobiology 64, 251–267. https://doi.org/10.1016/S0301-0082(00)00059-9

Scott, J.A., Da Camara, C.C., Early, J.E., 1999. Raloxifene: a selective estrogen receptor modulator. Am Fam Physician 60, 1131–1139.

St Pourcain, B., Robinson, E.B., Anttila, V., Sullivan, B.B., Maller, J., Golding, J., Skuse, D., Ring, S., Evans, D.M., Zammit, S., Fisher, S.E., Neale, B.M., Anney, R.J.L., Ripke, S., Hollegaard, M.V., Werge, T., Ronald, A., Grove, J., Hougaard, D.M., Børglum, A.D., Mortensen, P.B., Daly, M.J., Davey Smith, G., 2018. ASD and schizophrenia show distinct developmental profiles in common genetic overlap with populationbased social communication difficulties. Mol Psychiatry 23, 263–270. https://doi.org/10.1038/mp.2016.198

Stępnicki, P., Kondej, M., Kaczor, A.A., 2018. Current Concepts and Treatments of Schizophrenia. Molecules 23, 2087. https://doi.org/10.3390/molecules23082087

STRING v11: protein–protein association networks with increased coverage, supporting functional discovery in genome-wide experimental datasets | Nucleic Acids Research | Oxford Academic [WWW Document], n.d. URL https://academic.oup.com/nar/article/47/D1/D607/5198476 (accessed 9.28.21).

Stüve, O., Warnke, C., Deason, K., Stangel, M., Kieseier, B.C., Hartung, H.-P., von Büdingen, H.-C., Centonze, D., Forsthuber, T.G., Kappertz, V., 2014. CD19 as a molecular target in CNS autoimmunity. Acta Neuropathol 128, 177–190. https://doi.org/10.1007/s00401-014-1313-z

Sundermann, E.E., Maki, P.M., Bishop, J.R., 2010. A Review of Estrogen Receptor α Gene (ESR1) Polymorphisms, Mood, and Cognition. Menopause 17, 874–886. https://doi.org/10.1097/gme.0b013e3181df4a19

Walton, C., King, R., Rechtman, L., Kaye, W., Leray, E., Marrie, R.A., Robertson, N., La Rocca, N., Uitdehaag, B., van der Mei, I., Wallin, M., Helme, A., Angood Napier, C., Rijke, N., Baneke, P., 2020. Rising prevalence of multiple sclerosis worldwide: Insights from the Atlas of MS, third edition. Mult Scler 26, 1816–1821. https://doi.org/10.1177/1352458520970841

