## Supplementary material for "Identifying potential key genes and existing drugs for Multiple sclerosis, Schizophrenia, and Autism- an in silico approach": Gene mania report

### GeneMANIA report

Created on : 28 September 2021 23:08:47
Last database update : 13 August 2021 00:00:00
Application version : 3.6.0

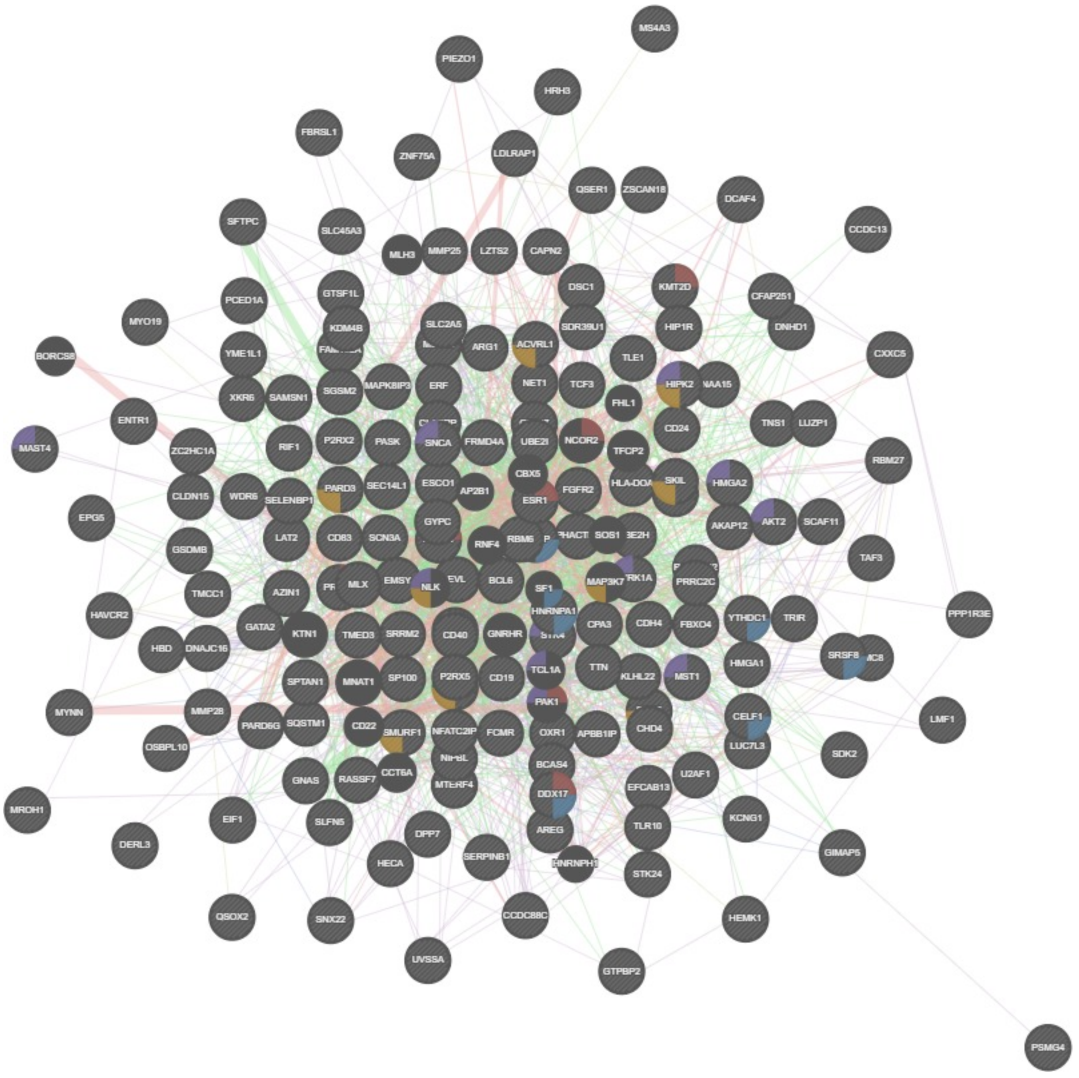


#### Networks

- Physical Interactions
- Co-expression
- Genetic Interactions
- Co-localization
- Shared protein domains
- Predicted
- Pathway

#### Functions

- intracellular steroid hormone receptor signaling pathway
- regulation of mRNA splicing, via spliceosome
- response to transforming growth factor beta
- peptidyl-serine modification

### Search parameters

**Organism** Homo sapiens (human)

**Genes** RBM6 , MMP25 , CCDC88C , SERPINB1 , KCNG1 , QSER1 , HIPK2 , FGFR2 , SP100 , MAST4 , SDK2 , TCF3 , GSDMB , APBB1IP , TNS1 , RIF1 , P2RX5 , YTHDC1 , MYNN , LAT2 , GNAS , RBM27 , ESR1 , CIRBP , RASSF7 , KLHL22 , DERL3 , DDX17 , SDR39U1 , CD40 , STK4 , HRH3 , STK24 , LMF1 , UBE2I , PIEZO1 , ZC2HC1A , CLASRP , AKT2 , ERF , CLDN15 , LZTS2 , MLX , LUC7L3 , AREG , SOX6 , CHD4 , PHACTR1 , CD83 , HECA , FBRSL1 , BCL6 , HEMK1 , FOXP1 , INO80D , PASK , DNAJC16 , RALGPS2 , PRRC2C , ARG1 , DCAF4 , ZSCAN18 , MTERF4 , GTSF1L , BCAS4 , KDM4B , SEC14L1 , HIP1R , AKAP12 , PCED1A , DSC1 , HAVCR2 , HNRNPA1 , SKIL , GYPC , YME1L1 , HMGA1 , MAPK8IP3 , SCAF11 , ACVRL1 , SGSM2 , ESCO1 , SLC2A5 , SELENBP1 , OSBPL10 , SNCA , PARD3 , CELF1 , MS4A3 , MS4A2 , HMGA2 , FRMD4A , FBXO4 , EPG5 , SCN3A , AZIN1 , SAMSN1 , TTN , DYRK1A , SNX22 , SKI , LDLRAP1 , EMSY , SLC45A3 , U2AF1 , PRMT2 , SQSTM1 , ZNF75A , FCMR , CAPN2 , CPA3 , UVSSA , NAA15 , NIPBL , OXR1 , TAF3 , QSOX2 , TMED3 , SLFN5 , FAM102A , KMT2D , TMC8 , SRRM2 , SFTPC , LUZP1 , XKR6 , CXXC5 , GTPBP2 , TMCC1 , MST1 , EIF1 , NET1 , TLR10 , NFATC2IP , DPP7 , CD19 , PARD6G , WDR6 , EFCAB13 , CDH4 , GATA2 , DNHD1 , MROH1 , PSMG4 , UBE2H , P2RX2 , GIMAP5 , EVL , TLE1 , SPTAN1 , SMURF1 , CD247 , HLA-DOA , HBD , PPP1R3E , CCDC13 , HBB , SRSF8 , MMP28 , CD24 , MYO19 , C19orf43 , WDR66 , SDCCAG3

**Network weighting** Automatically selected weighting method

**Networks**

**A**

Abbasi-Schild-Poulter-2019 , Abu-Odeh-Aqeilan-2014 , Achuthankutty-Mailand-2019 , Agrawal-Sedivy-2010 , Ahn-Lee-2008 , Albers-Koegl-2005 , Alexander-Wang-2018 , Alexandru-Deshaies-2008 , Alizadeh-Staudt-2000 , Alsulami-Cagney-2019 , An-Sun-2017 , Andresen-Flores-Morales-2014 , Arbogast-Gros-2019 , Arijs-Rutgeerts-2009 , Arroyo-Aloy-2014 , Arroyo-Aloy-2015 , Asadi-Dhanvantari-2018

**B**

Bailey-Hieter-2015 , Bandyopadhyay-Ideker-2010 , Banks-Washburn-2016 , Bantscheff-Drewes-2011 , Barr-Knapp-2009 , Barreiro-Alonso-Cerdán-2018 , Barrios-Rodiles-Wrana-2005 , Behrends-Harper-2010 , Behzadnia-Lührmann-2007 , Benleulmi-Chaachoua-Jockers-2016 A , Benleulmi-Chaachoua-Jockers-2016 B , Bennett-Harper-2010 , Benzinger-Hermeking-2005 , Berggård-James-2006 , Bett-Hay-2013 , Beyer-Boldt-2018 , Bhatnagar-Attie-2014 , Bild-Nevins-2006 B , BIOGRID-SMALL-SCALE-STUDIES , BIOGRID-SMALL-SCALE-STUDIES , Bishof-Seyfried-2018 , Blandin-Richard-2013 , Blomen-Brummelkamp-2015 ,

## B

Blomen-Brummelkamp-2015 , Bogachek-Weigel-2014 , Boldrick-Relman-2002 , Boldt-Roepman-2016 , Botham-Schimmer-2019 , Bouwmeester-Superti-Furga-2004 , Brady-Omary-2018 , Brajenovic-Drewes-2004 , Brehme-Superti-Furga-2009 , Burington-Shaughnessy-2008 , Butland-Hayden-2014 , Byron-Humphries-2012

## C

Cai-Conaway-2007 , Camargo-Brandon-2007 , Campos-Reinberg-2015 , Cao-Chinnaiyan-2014 , Carmon-Liu-2014 , Caron-van Attikum-2019 , CELL_MAP , Chen-Brown-2002 , Chen-Ge-2013 A , Chen-Ge-2013 B , Chen-Guan-2018 , Chen-Huang-2014 , Chen-Krogan-2018 , Chen-Yu-2018 , Chen-Zhang-2013 , Chen-Zhou-2019 , Cheng-DeCaprio-2017 , Chi-Reed-2018 , Chitale-Richly-2017 , Choi-Beutler-2019 , Choi-Busino-2018 , Choudhury-Michlewski-2017 , Christianson-Kopito-2011 , Cloutier-Coulombe-2013 , Cloutier-Coulombe-2017 , Colicelli-2010 , Colland-Gauthier-2004 , Conte-Perez-Oliva-2018 , Cooper-Green-2015 , Corominas-Iakoucheva-2014 , Couzens-Gingras-2013 , Cox-Rizzino-2013 , Coyaud-Raught-2015 , Crow-Cristea-2017

## D

Daakour-Twizere-2016 , Dabbaghizadeh-Tanguay-2018 , Dart-Wells-2015 , Das-Broemer-2019 , Davis-Glaunsinger-2015 , de Hoog-Mann-2004 , Devarajan-Ketha-Kumar-2012 , Diner-Cristea-2015 , Dittmer-Misteli-2014 , Dobbin-Giordano-2005 , Douanne-Bidère-2019 , Drissi-Boisvert-2015 , Du-Krogan-2017

## E

Elliott-Gyrd-Hansen-2016 , Emdal-Olsen-2015 , Enzo-Dupont-2015 , Ertych-Bastians-2016 , Ewing-Figeys-2007

## F

Fang-Lin-2011 , Faust-Frankel-2018 , Fenner-Prehn-2010 , Floyd-Pagliarini-2016 , Foerster-Ritter-2013 , Fogeron-Lange-2013 , Fonseca-Damgaard-2015 , Foster-Marshall-2013 , Fragoza-Yu-2019 , Freibaum-Taylor-2010

## G

Gabriel-Baumgrass-2016 , Gallardo-Vara-Bernabeu-2019 , Galligan-Howley-2015 , Gao-Reinberg-2012 , Gao-Vaziri-2016 , Garzia-Sonenberg-2017 , Gautier-Hall-2009 , Giannone-Liu-2010 , Gilmore-Washburn-2016 , Giurato-Tarallo-2018 , Glatter-Gstaiger-2009 , Gloeckner-Ueffing-2007 , Goehler-Wanker-2004 , Gordon-Krogan-2020 , Goudreault-Gingras-2009 , Greco-Cristea-2011 , Grossmann-Stelzl-2015 , Guarani-Harper-2014 , Guard-Old-2019 , Guardia-Laguarta-Przedborski-2019 , Guderian-Grimmler-2011 , Gupta-Pelletier-2015

## H

Han-Bassik-2017 A , Han-Bassik-2017 B , Hanson-Clayton-2014 , Hauri-Beisel-2016 , Hauri-Gstaiger-2013 , Havrylov-Redowicz-2009 , Havugimana-Emili-2012 , Hayes-Urbé-2012 , Hegele-Stelzl-2012 A , Hegele-Stelzl-2012 B , Heidelberger-Beli-2018 ,

## H

Hein-Mann-2015 , Hermjakob-Apweiler-2004 , Herr-Helleday-2015 , Hoffmeister-Längst-2017 , Horlbeck-Gilbert-2018 A , Horlbeck-Gilbert-2018 B , Hosp-Selbach-2015 , Hou-Chen-2018 , Hou-Huang-2017 , Hu-Woods-2019 , Hu-Yin-2019 , Hubel-Pichlmair-2019 , Huber-Hoelz-2017 , HUMANCYC , Humphries-Humphries-2009 , Hussain-Aldaz-2018 , Hutchins-Peters-2010 , Huttlin-Gygi-2015 , Huttlin-Harper-2017 , Hüttenhain-Krogan-2019

## I

I2D-BIND-Fly2Human , I2D-BIND-Mouse2Human , I2D-BIND-Rat2Human , I2D-BIND-Worm2Human , I2D-BIND-Yeast2Human , I2D-BioGRID-Fly2Human , I2D-BioGRID-Mouse2Human , I2D-BioGRID-Rat2Human , I2D-BioGRID-Worm2Human , I2D-BioGRID-Yeast2Human , I2D-Chen-Pawson-2009-PiwiScreen-Mouse2Human , I2D-Formstecher-Daviet-2005-Embryo-Fly2Human , I2D-Formstecher-Daviet-2005-Head-Fly2Human , I2D-Giot-Rothbert-2003-High-Fly2Human , I2D-Giot-Rothbert-2003-Low-Fly2Human , I2D-INNATEDB-Mouse2Human , I2D-IntAct-Fly2Human , I2D-IntAct-Mouse2Human , I2D-IntAct-Rat2Human , I2D-IntAct-Worm2Human , I2D-IntAct-Yeast2Human , I2D-Krogan-Greenblatt-2006-Core-Yeast2Human , I2D-Krogan-Greenblatt-2006-NonCore-Yeast2Human , I2D-Li-Vidal-2004-CE-DATA-Worm2Human , I2D-Li-Vidal-2004-CORE-1-Worm2Human , I2D-Li-Vidal-2004-CORE-2-Worm2Human , I2D-Li-Vidal-2004-interolog-Worm2Human , I2D-Li-Vidal-2004-literature-Worm2Human , I2D-Li-Vidal-2004-non-core-Worm2Human , I2D-Manual-Mouse2Human , I2D-Manual-Rat2Human , I2D-MGI-Mouse2Human , I2D-MINT-Fly2Human , I2D-MINT-Mouse2Human , I2D-MINT-Rat2Human , I2D-MINT-Worm2Human , I2D-MINT-Yeast2Human , I2D-MIPS-Yeast2Human , I2D-Ptacek-Snyder-2005-Yeast2Human , I2D-Stanyon-Finley-2004-CellCycle-Fly2Human , I2D-Tarassov-PCA-Yeast2Human , I2D-Tewari-Vidal-2004-TGFb-Worm2Human , I2D-vonMering-Bork-2002-High-Yeast2Human , I2D-vonMering-Bork-2002-Low-Yeast2Human , I2D-vonMering-Bork-2002-Medium-Yeast2Human , I2D-Wang-Orkin-2006-EScmplx-Mouse2Human , I2D-Wang-Orkin-2006-EScmplxIP-Mouse2Human , I2D-Wang-Orkin-2006-EScmplxlow-Mouse2Human , I2D-Yu-Vidal-2008-GoldStd-Yeast2Human , IMID , Ingham-Pawson-2005 , Innocenti-Brown-2011 , INTERPRO , Iradi-Borchelt-2018 , IREF-bhf-ucl , IREF-bind , IREF-bind-translation , IREF-biogrid , IREF-corum , IREF-dip , IREF-hpidb , IREF-hprd , IREF-huri , IREF-innatedb , IREF-intact , IREF-intcomplex , IREF-matrixdb , IREF-mbinfo , IREF-mint , IREF-mppi , IREF-quickgo , IREF-reactome , IREF-SMALL-SCALE-STUDIES , IREF-SMALL-SCALE-STUDIES , IREF-spike , IREF-uniprotpp , IREF-virushost , Ivanochko-Arrowsmith-2019

## J

Jain-Parker-2016 , Jang-Trono-2018 , Jeronimo-Coulombe-2007 , Jiang-de Kok-2017 , Jin-Pawson-2004 , Jirawatnotai-Sicinski-2011 , Johnson-Kerner-Wichterle-2015 , Johnson-Shoemaker-2003 , Jones-MacBeath-2006 , Joshi-Cristea-2013 , Jozwik-Carroll-2016 , Jäger-Krogan-2011

**K**

Kahle-Zoghbi-2011 , Kaltenbach-Hughes-2007 , Kang-Shin-2015 , Karras-Soengas-2019 , Kato-Sternberg-2014 , Katsogiannou-Rocchi-2014 , Kawahara-Paes Leme-2017 , Keller-Lee-2014 , Kennedy-Kolch-2020 A , Kennedy-Kolch-2020 B , Khanna-Parnaik-2018 , Kim-Major-2015 , Kneissl-Grummt-2003 , Koch-Hermeking-2007 , Kotlyar-Jurisica-2015 , Kristensen-Foster-2012 , Kumar-Maddika-2017 , Kumar-Vertegaal-2017 , Kupka-Walczak-2016 , Kärblane-Sarmiento-2015 , Kırlı-Görlich-2015

**L**

Lambert-Gingras-2015 , Lampert-Peter-2018 , Lau-Ronai-2012 , Lee-Choi-2016 , Lee-Choi-2017 , Lee-Jeong-2017 , Lee-Jou-2019 , Lee-Mayr-2019 , Lee-Songyang-2011 , Lehner-Sanderson-2004 A , Lehner-Sanderson-2004 B , Leung-Jones-2014 , Leung-Miller-2017 , Li-Chen-2015 , Li-Dorf-2011 A , Li-Dorf-2011 B , Li-Dorf-2014 , Li-Fu-2017 , Li-Haura-2013 , Li-Hung-2019 , Li-Lu-2018 , Li-Wang-2016 , Li-Zhou-2017 , Liebelt-Vertegaal-2020 , Lim-Zoghbi-2006 , Lin-Smith-2010 , Lipp-Guthrie-2015 , Liu-Chen-2019 , Liu-Sun-2019 , Liu-Takahashi-2017 , Liu-Tan-2018 , Liu-Varjosalo-2018 , Liu-Wang-2012 , Liu-Xu-2018 , Liu-Yang-2019 , Llères-Lamond-2010 , Loch-Strickler-2012 , Low-Heck-2014 , Lu-Bohr-2017 , Lu-Zhang-2013 , Luck-Calderwood-2020 , Lum-Cristea-2018 , Luo-Elledge-2009

**M**

Mak-Moffat-2010 , Malinová-Verheggen-2017 , Mallon-McKay-2013 , Malovannaya-Qin-2010 , Malty-Babu-2017 , Markson-Sanderson-2009 , Martin-Elledge-2017 , Maréchal-Zou-2014 , Matsumoto-Nakayama-2005 , Matsuoka-Elledge-2007 , McCracken-Blencowe-2005 , McFarland-Nussbaum-2008 , McNamara-D'Orso-2016 , Meek-Piwnica-Worms-2004 , Menon-Litovchick-2019 , Milev-Mouland-2012 , Miyamoto-Sato-Yanagawa-2010 , Mohammed-Carroll-2013 , Moon-Kim-2014 , Moutaoufik-Babu-2019 , Mugabo-Lim-2018 , Muller-Demeret-2012 , Murakawa-Landthaler-2015

**N**

Nakamura-Groth-2019 , Nakayama-Ohara-2002 , Napolitano-Meroni-2011 , Narayan-Bennett-2012 , Nassa-Weisz-2019 , Nathan-Goldberg-2013 , NCI_NATURE , Neganova-Lako-2011 , Newman-Keating-2003 , Noguchi-Kawahara-2018 , Nowak-Sommer-2019

**O**

Oliviero-Cagney-2015 , Oliviero-Cagney-2016 , Olma-Pintard-2009 , Oláh-Ovádi-2011 , Ouyang-Gill-2009

**P**

Panigrahi-Pati-2012 , Pankow-Yates-2015 , Pao-Virdee-2018 , Papp-Lamia-2015 , Pech-Settleman-2019 , Perez-Hernandez-Yáñez-Mó-2013 , Perez-Perri-Espinosa-2016 , Perou-Botstein-1999 , Perou-Botstein-2000 , Persaud-Rotin-2009 A , Persaud-Rotin-2009 B , Petschnigg-Stagljar-2014 , PFAM , Phillips-Corn-2013 , Pichlmair-

**P**

Superti-Furga-2011 , Pichlmair-Superti-Furga-2012 , Pilling-Cooper-2017 , Pladevall-Morera-Lopez-Contreras-2019 , Ptushkina-Ray-2017

**R**

Raisner-Gascoigne-2018 , Ramachandran-LaBaer-2004 , Raman-Harper-2015 , Ramaswamy-Golub-2001 , Ravasi-Hayashizaki-2010 , REACTOME , Reinke-Keating-2010 , Reinke-Keating-2013 , Rengasamy-Walsh-2017 , Reyniers-Taymans-2014 , Richter-Chrzanowska-Lightowlers-2010 , Rieger-Chu-2004 , Rivera-Paes Leme-2018 , Rodriguez-von Kriegsheim-2016 , Roewenstrunk-de la Luna-2019 , Rolland-Vidal-2014 , Rosenbluh-Hahn-2016 , Rosenwald-Staudt-2001 , Ross-Perou-2001 , Roth-Zlotnik-2006 , Rowbotham-Mermoud-2011 , Roy-Pardo-2014 , Roy-Parent-2013 , Rual-Vidal-2005

**S**

Saez-Vilchez-2018 , Sahni-Vidal-2015 , Saito-Kobarg-2017 , Sala-Ampe-2017 , Salvetti-Greco-2016 , Sang-Jackson-2011 , Sato-Conaway-2004 , Savidis-Brass-2016 , Schadt-Shoemaker-2004 , Schiza-Diamandis-2018 , Scholz-Taylor-2016 , Scifo-Lalowski-2015 , Scott-Guy-2017 , Scott-Schulman-2016 , Shami Shah-Baskin-2019 , Shen-Chen-2019 , Shen-Mali-2017 , Sherman-Teitell-2010 , Simabuco-Zanchin-2019 , Singh-Moore-2012 , So-Colwill-2015 , Sokolina-Stagljar-2017 , Soler-López-Aloy-2011 , Sowa-Harper-2009 , Srivas-Ideker-2016 , St-Denis-Gingras-2015 , St-Denis-Gingras-2016 , Stehling-Lill-2012 , Stehling-Lill-2013 , Stelzl-Wanker-2005 , Stuart-Kim-2003 , Sundell-Ivarsson-2018 , Suter-Wanker-2013 , Swayampakula-Dedhar-2017

**T**

Taipale-Lindquist-2012 , Taipale-Lindquist-2014 , Takahashi-Conaway-2011 , Tang-Wang-2019 , Tarallo-Weisz-2011 , Teixeira-Gomes-2010 , Teixeira-Laman-2016 A , Teixeira-Laman-2016 B , Thalappilly-Dusetti-2008 , Thompson-Luchansky-2014 , Tiemann-Kani-2019 , Tomkins-Manzoni-2018 , Tong-Moran-2014 , Toyoshima-Grandori-2012 , Trepte-Wanker-2018 A , Trepte-Wanker-2018 B , Tsai-Cristea-2012

**U**

Ugidos-Vandenbroeck-2019

**V**

Van Acker-Dewilde-2019 , Van Alstyne-Pellizzoni-2018 , Van Quickelberghe-Gevaert-2018 , van Wijk-Timmers-2009 , Vandamme-Angrand-2011 , Varier-Vermeulen-2016 , Varjosalo-Gstaiger-2013 A , Varjosalo-Gstaiger-2013 B , Varjosalo-Superti-Furga-2013 , Vastrik-Stein-2007 , Venkatesan-Vidal-2009 , Viita-Vartiainen-2019 , Vinayagam-Wanker-2011 , Virok-Fülöp-2011 , Vizeacoumar-Moffat-2013 , von Hundelshausen-Weber-2017

**W**

Wallach-Kramer-2013 , Wan-Emili-2015 , Wang-Balch-2006 , Wang-Cheung-2015 ,

**W**

Wang-He-2008 , Wang-Huang-2017 , Wang-Liu-2019 , Wang-Maris-2006 , Wang-Xiong-2019 , Wang-Xu-2015 , Wang-Yang-2011 , Watanabe-Fujita-2018 , Weimann-Stelzl-2013 A , Weimann-Stelzl-2013 B , Weinmann-Meister-2009 , Weishäupl-Schmidt-2019 , Weith-Meyer-2018 , Whisenant-Salomon-2015 , Wilkinson-Coba-2019 , Willingham-Muchowski-2003 , Winczura-Jensen-2018 , Wong-O'Bryan-2012 , Woods-Monteiro-2012 A , Woods-Monteiro-2012 B , Woodsmith-Sanderson-2012 , Wu-Garvey-2007 , Wu-Li-2007 , Wu-Ma-2012 , Wu-Stein-2010 , Wu-Stein-2010

**X**

Xiao-Brown-2018 , Xiao-Lefkowitz-2007 , Xie-Cong-2013 , Xie-Green-2012 , Xie-Zhang-2017 , Xu-Ye-2012 , Xu-Zetter-2016

**Y**

Yachie-Roth-2016 , Yadav-Varjosalo-2017 , Yamauchi-Maeda-2018 , Yang-Brasier-2015 , Yang-Chen-2010 , Yang-Maurer-2018 , Yang-Vidal-2016 , Yang-Wang-2018 , Yao-Stagljar-2017 A , Yao-Stagljar-2017 B , Yatim-Benkirane-2012 , Yeung-Dougan-2019 , Yu-Chow-2013 , Yu-Engel-2018 , Yu-Vidal-2011 , Yue-Liu-2018

**Z**

Zanon-Pichler-2013 , Zeller-Wei-2006 , Zhang-Shang-2006 , Zhang-Vermeulen-2017 , Zhang-Wang-2018 , Zhang-Wheeler-2014 , Zhang-Xu-2018 , Zhang-Zou-2011 , Zhao-Krug-2005 , Zhao-Yang-2011 , Zhong-Vidal-2016 , Zhou-Conrads-2004 , Zhou-Hanemann-2016 , Zhu-Liu-2018

### Genes

| Gene | Description | Rank |
|---|---|---|
| MS4A3 | membrane spanning 4-domains A3 [Source:HGNC Symbol;Acc:HGNC:7317] | N/A |
| PSMG4 | proteasome assembly chaperone 4 [Source:HGNC Symbol;Acc:HGNC:21108] | N/A |
| CCDC13 | coiled-coil domain containing 13 [Source:HGNC Symbol;Acc:HGNC:26358] | N/A |
| MROH1 | maestro heat like repeat family member 1 [Source:HGNC Symbol;Acc:HGNC:26958] | N/A |
| XKR6 | XK related 6 [Source:HGNC Symbol;Acc:HGNC:27806] | N/A |
| QSOX2 | quiescin sulfhydryl oxidase 2 [Source:HGNC Symbol;Acc:HGNC:30249] | N/A |
| DNHD1 | dynein heavy chain domain 1 [Source:HGNC Symbol;Acc:HGNC:26532] | N/A |
| PPP1R3E | protein phosphatase 1 regulatory subunit 3E [Source:HGNC Symbol;Acc:HGNC:14943] | N/A |
| DERL3 | derlin 3 [Source:HGNC Symbol;Acc:HGNC:14236] | N/A |
| ZNF75A | zinc finger protein 75a [Source:HGNC Symbol;Acc:HGNC:13146] | N/A |
| EFCAB13 | EF-hand calcium binding domain 13 [Source:HGNC Symbol;Acc:HGNC:26864] | N/A |
| CFAP251 | cilia and flagella associated protein 251 [Source:HGNC Symbol;Acc:HGNC:28506] | N/A |
| SLFN5 | schlafen family member 5 [Source:HGNC Symbol;Acc:HGNC:28286] | N/A |
| TMC8 | transmembrane channel like 8 [Source:HGNC Symbol;Acc:HGNC:20474] | N/A |
| GTSF1L | gametocyte specific factor 1 like [Source:HGNC Symbol;Acc:HGNC:16198] | N/A |
| TAF3 | TATA-box binding protein associated factor 3 [Source:HGNC Symbol;Acc:HGNC:17303] | N/A |
| SDK2 | sidekick cell adhesion molecule 2 [Source:HGNC Symbol;Acc:HGNC:19308] | N/A |
| LMF1 | lipase maturation factor 1 [Source:HGNC Symbol;Acc:HGNC:14154] | N/A |
| PCED1A | PC-esterase domain containing 1A [Source:HGNC Symbol;Acc:HGNC:16212] | N/A |
| EPG5 | ectopic P-granules autophagy protein 5 homolog [Source:HGNC Symbol;Acc:HGNC:29331] | N/A |
| HAVCR2 | hepatitis A virus cellular receptor 2 [Source:HGNC Symbol;Acc:HGNC:18437] | N/A |

| Gene | Description | Rank |
|---|---|---|
| DPP7 | dipeptidyl peptidase 7 [Source:HGNC Symbol;Acc:HGNC:14892] | N/A |
| SLC45A3 | solute carrier family 45 member 3 [Source:HGNC Symbol;Acc:HGNC:8642] | N/A |
| P2RX2 | purinergic receptor P2X 2 [Source:HGNC Symbol;Acc:HGNC:15459] | N/A |
| LZTS2 | leucine zipper tumor suppressor 2 [Source:HGNC Symbol;Acc:HGNC:29381] | N/A |
| LAT2 | linker for activation of T cells family member 2 [Source:HGNC Symbol;Acc:HGNC:12749] | N/A |
| ZSCAN18 | zinc finger and SCAN domain containing 18 [Source:HGNC Symbol;Acc:HGNC:21037] | N/A |
| MMP28 | matrix metallopeptidase 28 [Source:HGNC Symbol;Acc:HGNC:14366] | N/A |
| SCN3A | sodium voltage-gated channel alpha subunit 3 [Source:HGNC Symbol;Acc:HGNC:10590] | N/A |
| HRH3 | histamine receptor H3 [Source:HGNC Symbol;Acc:HGNC:5184] | N/A |
| PARD6G | par-6 family cell polarity regulator gamma [Source:HGNC Symbol;Acc:HGNC:16076] | N/A |
| OXR1 | oxidation resistance 1 [Source:HGNC Symbol;Acc:HGNC:15822] | N/A |
| SNX22 | sorting nexin 22 [Source:HGNC Symbol;Acc:HGNC:16315] | N/A |
| HEMK1 | HemK methyltransferase family member 1 [Source:HGNC Symbol;Acc:HGNC:24923] | N/A |
| TRIR | telomerase RNA component interacting RNase [Source:HGNC Symbol;Acc:HGNC:28424] | N/A |
| INO80D | INO80 complex subunit D [Source:HGNC Symbol;Acc:HGNC:25997] | N/A |
| ENTR1 | endosome associated trafficking regulator 1 [Source:HGNC Symbol;Acc:HGNC:10667] | N/A |
| CCDC88C | coiled-coil domain containing 88C [Source:HGNC Symbol;Acc:HGNC:19967] | N/A |
| MYO19 | myosin XIX [Source:HGNC Symbol;Acc:HGNC:26234] | N/A |
| LDLRAP1 | low density lipoprotein receptor adaptor protein 1 [Source:HGNC Symbol;Acc:HGNC:18640] | N/A |
| WDR6 | WD repeat domain 6 [Source:HGNC Symbol;Acc:HGNC:12758] | N/A |
| SOX6 | SRY-box transcription factor 6 [Source:HGNC Symbol;Acc:HGNC:16421] | N/A |
| UVSSA | UV stimulated scaffold protein A [Source:HGNC Symbol;Acc:HGNC:29304] | N/A |
| MAST4 | microtubule associated serine/threonine kinase family member 4 [Source:HGNC Symbol;Acc:HGNC:19037] | N/A |

| Gene | Description | Rank |
|---|---|---|
| FBRSL1 | fibrosin like 1 [Source:HGNC Symbol;Acc:HGNC:29308] | N/A |
| MYNN | myoneurin [Source:HGNC Symbol;Acc:HGNC:14955] | N/A |
| CXXC5 | CXXC finger protein 5 [Source:HGNC Symbol;Acc:HGNC:26943] | N/A |
| PHACTR1 | phosphatase and actin regulator 1 [Source:HGNC Symbol;Acc:HGNC:20990] | N/A |
| YME1L1 | YME1 like 1 ATPase [Source:HGNC Symbol;Acc:HGNC:12843] | N/A |
| TMCC1 | transmembrane and coiled-coil domain family 1 [Source:HGNC Symbol;Acc:HGNC:29116] | N/A |
| MTERF4 | mitochondrial transcription termination factor 4 [Source:HGNC Symbol;Acc:HGNC:28785] | N/A |
| GSDMB | gasdermin B [Source:HGNC Symbol;Acc:HGNC:23690] | N/A |
| RBM27 | RNA binding motif protein 27 [Source:HGNC Symbol;Acc:HGNC:29243] | N/A |
| GATA2 | GATA binding protein 2 [Source:HGNC Symbol;Acc:HGNC:4171] | N/A |
| FBXO4 | F-box protein 4 [Source:HGNC Symbol;Acc:HGNC:13583] | N/A |
| TNS1 | tensin 1 [Source:HGNC Symbol;Acc:HGNC:11973] | N/A |
| ZC2HC1A | zinc finger C2HC-type containing 1A [Source:HGNC Symbol;Acc:HGNC:24277] | N/A |
| QSER1 | glutamine and serine rich 1 [Source:HGNC Symbol;Acc:HGNC:26154] | N/A |
| FRMD4A | FERM domain containing 4A [Source:HGNC Symbol;Acc:HGNC:25491] | N/A |
| KMT2D | lysine methyltransferase 2D [Source:HGNC Symbol;Acc:HGNC:7133] | N/A |
| RIF1 | replication timing regulatory factor 1 [Source:HGNC Symbol;Acc:HGNC:23207] | N/A |
| HECA | hdc homolog, cell cycle regulator [Source:HGNC Symbol;Acc:HGNC:21041] | N/A |
| ESCO1 | establishment of sister chromatid cohesion N-acetyltransferase 1 [Source:HGNC Symbol;Acc:HGNC:24645] | N/A |
| RALGPS2 | Ral GEF with PH domain and SH3 binding motif 2 [Source:HGNC Symbol;Acc:HGNC:30279] | N/A |
| BCAS4 | breast carcinoma amplified sequence 4 [Source:HGNC Symbol;Acc:HGNC:14367] | N/A |
| SFTPC | surfactant protein C [Source:HGNC Symbol;Acc:HGNC:10802] | N/A |
| FAM102A | family with sequence similarity 102 member A [Source:HGNC Symbol;Acc:HGNC:31419] | N/A |
| LUZP1 | leucine zipper protein 1 [Source:HGNC Symbol;Acc:HGNC:14985] | N/A |
| GTPBP2 | GTP binding protein 2 [Source:HGNC Symbol;Acc:HGNC:4670] | N/A |

| Gene | Description | Rank |
|---|---|---|
| PARD3 | par-3 family cell polarity regulator [Source:HGNC Symbol;Acc:HGNC:16051] | N/A |
| EIF1 | eukaryotic translation initiation factor 1 [Source:HGNC Symbol;Acc:HGNC:3249] | N/A |
| KDM4B | lysine demethylase 4B [Source:HGNC Symbol;Acc:HGNC:29136] | N/A |
| EMSY | EMSY transcriptional repressor, BRCA2 interacting [Source:HGNC Symbol;Acc:HGNC:18071] | N/A |
| SRSF8 | serine and arginine rich splicing factor 8 [Source:HGNC Symbol;Acc:HGNC:16988] | N/A |
| GIMAP5 | GTPase, IMAP family member 5 [Source:HGNC Symbol;Acc:HGNC:18005] | N/A |
| KCNG1 | potassium voltage-gated channel modifier subfamily G member 1 [Source:HGNC Symbol;Acc:HGNC:6248] | N/A |
| PRMT2 | protein arginine methyltransferase 2 [Source:HGNC Symbol;Acc:HGNC:5186] | N/A |
| OSBPL10 | oxysterol binding protein like 10 [Source:HGNC Symbol;Acc:HGNC:16395] | N/A |
| YTHDC1 | YTH domain containing 1 [Source:HGNC Symbol;Acc:HGNC:30626] | N/A |
| MMP25 | matrix metallopeptidase 25 [Source:HGNC Symbol;Acc:HGNC:14246] | N/A |
| EVL | Enah/Vasp-like [Source:HGNC Symbol;Acc:HGNC:20234] | N/A |
| MS4A2 | membrane spanning 4-domains A2 [Source:HGNC Symbol;Acc:HGNC:7316] | N/A |
| HMGA2 | high mobility group AT-hook 2 [Source:HGNC Symbol;Acc:HGNC:5009] | N/A |
| MLX | MAX dimerization protein MLX [Source:HGNC Symbol;Acc:HGNC:11645] | N/A |
| KLHL22 | kelch like family member 22 [Source:HGNC Symbol;Acc:HGNC:25888] | N/A |
| HIP1R | huntingtin interacting protein 1 related [Source:HGNC Symbol;Acc:HGNC:18415] | N/A |
| CLDN15 | claudin 15 [Source:HGNC Symbol;Acc:HGNC:2036] | N/A |
| NAA15 | N-alpha-acetyltransferase 15, NatA auxiliary subunit [Source:HGNC Symbol;Acc:HGNC:30782] | N/A |
| CD24 | CD24 molecule [Source:HGNC Symbol;Acc:HGNC:1645] | N/A |
| TLR10 | toll like receptor 10 [Source:HGNC Symbol;Acc:HGNC:15634] | N/A |
| MAPK8IP3 | mitogen-activated protein kinase 8 interacting protein 3 [Source:HGNC Symbol;Acc:HGNC:6884] | N/A |
| PIEZO1 | piezo type mechanosensitive ion channel component 1 [Source:HGNC | N/A |

| Gene | Description | Rank |
|---|---|---|
| | Symbol;Acc:HGNC:28993] | |
| DNAJC16 | DnaJ heat shock protein family (Hsp40) member C16 [Source:HGNC Symbol;Acc:HGNC:29157] | N/A |
| HBD | hemoglobin subunit delta [Source:HGNC Symbol;Acc:HGNC:4829] | N/A |
| SKI | SKI proto-oncogene [Source:HGNC Symbol;Acc:HGNC:10896] | N/A |
| HIPK2 | homeodomain interacting protein kinase 2 [Source:HGNC Symbol;Acc:HGNC:14402] | N/A |
| AREG | amphiregulin [Source:HGNC Symbol;Acc:HGNC:651] | N/A |
| NIPBL | NIPBL cohesin loading factor [Source:HGNC Symbol;Acc:HGNC:28862] | N/A |
| TLE1 | TLE family member 1, transcriptional corepressor [Source:HGNC Symbol;Acc:HGNC:11837] | N/A |
| DSC1 | desmocollin 1 [Source:HGNC Symbol;Acc:HGNC:3035] | N/A |
| TMED3 | transmembrane p24 trafficking protein 3 [Source:HGNC Symbol;Acc:HGNC:28889] | N/A |
| APBB1IP | amyloid beta precursor protein binding family B member 1 interacting protein [Source:HGNC Symbol;Acc:HGNC:17379] | N/A |
| UBE2H | ubiquitin conjugating enzyme E2 H [Source:HGNC Symbol;Acc:HGNC:12484] | N/A |
| HLA-DOA | major histocompatibility complex, class II, DO alpha [Source:HGNC Symbol;Acc:HGNC:4936] | N/A |
| NET1 | neuroepithelial cell transforming 1 [Source:HGNC Symbol;Acc:HGNC:14592] | N/A |
| NFATC2IP | nuclear factor of activated T cells 2 interacting protein [Source:HGNC Symbol;Acc:HGNC:25906] | N/A |
| SKIL | SKI like proto-oncogene [Source:HGNC Symbol;Acc:HGNC:10897] | N/A |
| ERF | ETS2 repressor factor [Source:HGNC Symbol;Acc:HGNC:3444] | N/A |
| TCF3 | transcription factor 3 [Source:HGNC Symbol;Acc:HGNC:11633] | N/A |
| CD247 | CD247 molecule [Source:HGNC Symbol;Acc:HGNC:1677] | N/A |
| CPA3 | carboxypeptidase A3 [Source:HGNC Symbol;Acc:HGNC:2298] | N/A |
| SCAF11 | SR-related CTD associated factor 11 [Source:HGNC Symbol;Acc:HGNC:10784] | N/A |
| SLC2A5 | solute carrier family 2 member 5 [Source:HGNC Symbol;Acc:HGNC:11010] | N/A |
| SEC14L1 | SEC14 like lipid binding 1 [Source:HGNC Symbol;Acc:HGNC:10698] | N/A |
| LUC7L3 | LUC7 like 3 pre-mRNA splicing factor [Source:HGNC Symbol;Acc:HGNC:24309] | N/A |

| Gene | Description | Rank |
|---|---|---|
| PRRC2C | proline rich coiled-coil 2C [Source:HGNC Symbol;Acc:HGNC:24903] | N/A |
| DDX17 | DEAD-box helicase 17 [Source:HGNC Symbol;Acc:HGNC:2740] | N/A |
| SGSM2 | small G protein signaling modulator 2 [Source:HGNC Symbol;Acc:HGNC:29026] | N/A |
| CELF1 | CUGBP Elav-like family member 1 [Source:HGNC Symbol;Acc:HGNC:2549] | N/A |
| PASK | PAS domain containing serine/threonine kinase [Source:HGNC Symbol;Acc:HGNC:17270] | N/A |
| CDH4 | cadherin 4 [Source:HGNC Symbol;Acc:HGNC:1763] | N/A |
| AZIN1 | antizyme inhibitor 1 [Source:HGNC Symbol;Acc:HGNC:16432] | N/A |
| U2AF1 | U2 small nuclear RNA auxiliary factor 1 [Source:HGNC Symbol;Acc:HGNC:12453] | N/A |
| FOXP1 | forkhead box P1 [Source:HGNC Symbol;Acc:HGNC:3823] | N/A |
| AKT2 | AKT serine/threonine kinase 2 [Source:HGNC Symbol;Acc:HGNC:392] | N/A |
| SQSTM1 | sequestosome 1 [Source:HGNC Symbol;Acc:HGNC:11280] | N/A |
| STK24 | serine/threonine kinase 24 [Source:HGNC Symbol;Acc:HGNC:11403] | N/A |
| CAPN2 | calpain 2 [Source:HGNC Symbol;Acc:HGNC:1479] | N/A |
| MST1 | macrophage stimulating 1 [Source:HGNC Symbol;Acc:HGNC:7380] | N/A |
| HBB | hemoglobin subunit beta [Source:HGNC Symbol;Acc:HGNC:4827] | N/A |
| FGFR2 | fibroblast growth factor receptor 2 [Source:HGNC Symbol;Acc:HGNC:3689] | N/A |
| SDR39U1 | short chain dehydrogenase/reductase family 39U member 1 [Source:HGNC Symbol;Acc:HGNC:20275] | N/A |
| ARG1 | arginase 1 [Source:HGNC Symbol;Acc:HGNC:663] | N/A |
| SAMSN1 | SAM domain, SH3 domain and nuclear localization signals 1 [Source:HGNC Symbol;Acc:HGNC:10528] | N/A |
| CLASRP | CLK4 associating serine/arginine rich protein [Source:HGNC Symbol;Acc:HGNC:17731] | N/A |
| CIRBP | cold inducible RNA binding protein [Source:HGNC Symbol;Acc:HGNC:1982] | N/A |
| ACVRL1 | activin A receptor like type 1 [Source:HGNC Symbol;Acc:HGNC:175] | N/A |
| GNAS | GNAS complex locus [Source:HGNC Symbol;Acc:HGNC:4392] | N/A |
| CD40 | CD40 molecule [Source:HGNC Symbol;Acc:HGNC:11919] | N/A |
| RASSF7 | Ras association domain family member 7 [Source:HGNC Symbol;Acc:HGNC:1166] | N/A |

| Gene | Description | Rank |
|---|---|---|
| STK4 | serine/threonine kinase 4 [Source:HGNC Symbol;Acc:HGNC:11408] | N/A |
| AKAP12 | A-kinase anchoring protein 12 [Source:HGNC Symbol;Acc:HGNC:370] | N/A |
| SELENBP1 | selenium binding protein 1 [Source:HGNC Symbol;Acc:HGNC:10719] | N/A |
| RBM6 | RNA binding motif protein 6 [Source:HGNC Symbol;Acc:HGNC:9903] | N/A |
| SP100 | SP100 nuclear antigen [Source:HGNC Symbol;Acc:HGNC:11206] | N/A |
| DCAF4 | DDB1 and CUL4 associated factor 4 [Source:HGNC Symbol;Acc:HGNC: 20229] | N/A |
| SERPINB1 | serpin family B member 1 [Source:HGNC Symbol;Acc:HGNC:3311] | N/A |
| P2RX5 | purinergic receptor P2X 5 [Source:HGNC Symbol;Acc:HGNC:8536] | N/A |
| FCMR | Fc fragment of IgM receptor [Source:HGNC Symbol;Acc:HGNC:14315] | N/A |
| GYPC | glycophorin C (Gerbich blood group) [Source:HGNC Symbol;Acc:HGNC: 4704] | N/A |
| HMGA1 | high mobility group AT-hook 1 [Source:HGNC Symbol;Acc:HGNC:5010] | N/A |
| SPTAN1 | spectrin alpha, non-erythrocytic 1 [Source:HGNC Symbol;Acc:HGNC: 11273] | N/A |
| BCL6 | BCL6 transcription repressor [Source:HGNC Symbol;Acc:HGNC:1001] | N/A |
| CD83 | CD83 molecule [Source:HGNC Symbol;Acc:HGNC:1703] | N/A |
| SNCA | synuclein alpha [Source:HGNC Symbol;Acc:HGNC:11138] | N/A |
| CD19 | CD19 molecule [Source:HGNC Symbol;Acc:HGNC:1633] | N/A |
| TTN | titin [Source:HGNC Symbol;Acc:HGNC:12403] | N/A |
| SRRM2 | serine/arginine repetitive matrix 2 [Source:HGNC Symbol;Acc:HGNC: 16639] | N/A |
| UBE2I | ubiquitin conjugating enzyme E2 I [Source:HGNC Symbol;Acc:HGNC: 12485] | N/A |
| HNRNPA1 | heterogeneous nuclear ribonucleoprotein A1 [Source:HGNC Symbol;Acc: HGNC:5031] | N/A |
| SMURF1 | SMAD specific E3 ubiquitin protein ligase 1 [Source:HGNC Symbol;Acc: HGNC:16807] | N/A |
| DYRK1A | dual specificity tyrosine phosphorylation regulated kinase 1A [Source: HGNC Symbol;Acc:HGNC:3091] | N/A |
| CHD4 | chromodomain helicase DNA binding protein 4 [Source:HGNC Symbol; Acc:HGNC:1919] | N/A |
| ESR1 | estrogen receptor 1 [Source:HGNC Symbol;Acc:HGNC:3467] | N/A |
| MNAT1 | MNAT1 component of CDK activating kinase [Source:HGNC Symbol; Acc:HGNC:7181] | 1 |

| Gene | Description | Rank |
|---|---|---|
| KTN1 | kinectin 1 [Source:HGNC Symbol;Acc:HGNC:6467] | 2 |
| NCOR2 | nuclear receptor corepressor 2 [Source:HGNC Symbol;Acc:HGNC:7673] | 3 |
| GNRHR | gonadotropin releasing hormone receptor [Source:HGNC Symbol;Acc: HGNC:4421] | 4 |
| TFCP2 | transcription factor CP2 [Source:HGNC Symbol;Acc:HGNC:11748] | 5 |
| MLH3 | mutL homolog 3 [Source:HGNC Symbol;Acc:HGNC:7128] | 6 |
| PAK1 | p21 (RAC1) activated kinase 1 [Source:HGNC Symbol;Acc:HGNC:8590] | 7 |
| MAP3K7 | mitogen-activated protein kinase kinase kinase 7 [Source:HGNC Symbol; Acc:HGNC:6859] | 8 |
| CD22 | CD22 molecule [Source:HGNC Symbol;Acc:HGNC:1643] | 9 |
| CCT6A | chaperonin containing TCP1 subunit 6A [Source:HGNC Symbol;Acc: HGNC:1620] | 10 |
| CBX5 | chromobox 5 [Source:HGNC Symbol;Acc:HGNC:1555] | 11 |
| BORCS8 | BLOC-1 related complex subunit 8 [Source:HGNC Symbol;Acc:HGNC: 37247] | 12 |
| NLK | nemo like kinase [Source:HGNC Symbol;Acc:HGNC:29858] | 13 |
| RNF4 | ring finger protein 4 [Source:HGNC Symbol;Acc:HGNC:10067] | 14 |
| TCL1A | T cell leukemia/lymphoma 1A [Source:HGNC Symbol;Acc:HGNC:11648] | 15 |
| SOS1 | SOS Ras/Rac guanine nucleotide exchange factor 1 [Source:HGNC Symbol;Acc:HGNC:11187] | 16 |
| AP2B1 | adaptor related protein complex 2 subunit beta 1 [Source:HGNC Symbol; Acc:HGNC:563] | 17 |
| HNRNPH1 | heterogeneous nuclear ribonucleoprotein H1 [Source:HGNC Symbol;Acc: HGNC:5041] | 18 |
| SF1 | splicing factor 1 [Source:HGNC Symbol;Acc:HGNC:12950] | 19 |
| FHL1 | four and a half LIM domains 1 [Source:HGNC Symbol;Acc:HGNC:3702] | 20 |

### Networks

#### **Physical Interactions** 53.83%

##### Bandyopadhyay-Ideker-2010 5.09%

A human MAP kinase interactome. Bandyopadhyay et al (2010). *Nat Methods*

Physical Interactions with 653 interactions from iRefIndex

##### Jeronimo-Coulombe-2007 4.91%

Systematic analysis of the protein interaction network for the human transcription machinery reveals the identity of the 7SK capping enzyme. Jeronimo et al (2007). *Mol Cell*

Physical Interactions with 699 interactions from BioGRID

##### Nassa-Weisz-2019 4.24%

The RNA-mediated estrogen receptor   interactome of hormone-dependent human breast cancer cell nuclei. Nassa et al (2019). *Sci Data*

Physical Interactions with 1,490 interactions from BioGRID

##### Tarallo-Weisz-2011 3.89%

Identification of proteins associated with ligand-activated estrogen receptor   in human breast cancer cell nuclei by tandem affinity purification and nano LC-MS/MS. Tarallo et al (2011). *Proteomics*

Physical Interactions with 244 interactions from BioGRID

##### Xie-Cong-2013 3.62%

Deubiquitinase FAM/USP9X interacts with the E3 ubiquitin ligase SMURF1 protein and protects it from ligase activity-dependent self-degradation. Xie et al (2013). *J Biol Chem*

Physical Interactions with 170 interactions from BioGRID

##### Roewenstrunk-de la Luna-2019 3.47%

A comprehensive proteomics-based interaction screen that links DYRK1A to RNF169 and to the DNA damage response. Roewenstrunk et al (2019). *Sci Rep*

Physical Interactions with 116 interactions from BioGRID

##### Hoffmeister-Längst-2017 3.36%

CHD3 and CHD4 form distinct NuRD complexes with different yet overlapping functionality. Hoffmeister et al (2017). *Nucleic Acids Res*

Physical Interactions with 1,015 interactions from BioGRID

##### Blandin-Richard-2013 3.20%

A human skeletal muscle interactome centered on proteins involved in muscular dystrophies: LGMD interactome. Blandin et al (2013). *Skelet Muscle*

Physical Interactions with 655 interactions from iRefIndex

##### Mohammed-Carroll-2013 2.21%

Endogenous purification reveals GREB1 as a key estrogen receptor regulatory factor. Mohammed et al (2013). *Cell Rep*

Physical Interactions with 112 interactions from BioGRID

##### Liu-Yang-2019 2.07%

Inflammation-dependent overexpression of c-Myc enhances CRL4<sup>DCAF4</sup> E3 ligase activity and promotes ubiquitination of ST7 in colitis-associated cancer. Liu et al (2019). *J Pathol*

Physical Interactions with 279 interactions from BioGRID

##### Yachie-Roth-2016 2.04%

#### **Physical Interactions** 53.83%

##### Yachie-Roth-2016

Pooled-matrix protein interaction screens using Barcode Fusion Genetics. Yachie et al (2016). *Mol Syst Biol*

Physical Interactions with 671 interactions from BioGRID

##### Zhong-Vidal-2016 2.04%

An inter-species protein-protein interaction network across vast evolutionary distance. Zhong et al (2016). *Mol Syst Biol*

Physical Interactions with 472 interactions from BioGRID

##### McCracken-Blencowe-2005 1.84%

Proteomic analysis of SRm160-containing complexes reveals a conserved association with cohesin. McCracken et al (2005). *J Biol Chem*

Physical Interactions with 198 interactions from BioGRID

##### Sowa-Harper-2009 1.51%

Defining the human deubiquitinating enzyme interaction landscape. Sowa et al (2009). *Cell*

Physical Interactions with 1,509 interactions from BioGRID

##### Li-Chen-2015 1.48%

Proteomic analyses reveal distinct chromatin-associated and soluble transcription factor complexes. Li et al (2015). *Mol Syst Biol*

Physical Interactions with 1,811 interactions from BioGRID

##### Guard-Old-2019 1.46%

The nuclear interactome of DYRK1A reveals a functional role in DNA damage repair. Guard et al (2019). *Sci Rep*

Physical Interactions with 105 interactions from BioGRID

##### Varier-Vermeulen-2016 1.34%

Recruitment of the Mammalian Histone-modifying EMSY Complex to Target Genes Is Regulated by ZNF131. Varier et al (2016). *J Biol Chem*

Physical Interactions with 141 interactions from BioGRID

##### IREF-innatedb 1.00%

Physical Interactions with 2,355 interactions from iRefIndex

##### Boldt-Roepman-2016 0.84%

An organelle-specific protein landscape identifies novel diseases and molecular mechanisms. Boldt et al (2016). *Nat Commun*

Physical Interactions with 4,898 interactions from BioGRID

##### Varjosalo-Gstaiger-2013 A 0.74%

The protein interaction landscape of the human CMGC kinase group. Varjosalo et al (2013). *Cell Rep*

Physical Interactions with 690 interactions from BioGRID

##### IREF-bind-translation 0.53%

Physical Interactions with 6,056 interactions from iRefIndex

##### IREF-mint 0.43%

Physical Interactions with 14,408 interactions from iRefIndex

##### IREF-reactome 0.43%

Physical Interactions with 111,926 interactions from iRefIndex

##### Vastrik-Stein-2007 0.43%

Reactome: a knowledge base of biologic pathways and processes. Vastrik et al (2007). *Genome Biol*

#### Physical Interactions 53.83%

##### Vastrik-Stein-2007

Physical Interactions with 111,926 interactions from iRefIndex

##### IREF-hprd 0.36%

Physical Interactions with 33,375 interactions from iRefIndex

##### Karras-Soengas-2019 0.32%

p62/SQSTM1 Fuels Melanoma Progression by Opposing mRNA Decay of a Selective Set of Pro-metastatic Factors. Karras et al (2019). *Cancer Cell*

Physical Interactions with 147 interactions from BioGRID

##### McFarland-Nussbaum-2008 0.30%

Proteomics analysis identifies phosphorylation-dependent alpha-synuclein protein interactions. McFarland et al (2008). *Mol Cell Proteomics*

Physical Interactions with 159 interactions from BioGRID

##### Roy-Pardo-2014 0.17%

hnRNPA1 couples nuclear export and translation of specific mRNAs downstream of FGF-2/S6K2 signalling. Roy et al (2014). *Nucleic Acids Res*

Physical Interactions with 386 interactions from BioGRID

##### Menon-Litovchick-2019 0.14%

DYRK1A regulates the recruitment of 53BP1 to the sites of DNA damage in part through interaction with RNF169. Menon et al (2019). *Cell Cycle*

Physical Interactions with 119 interactions from BioGRID

##### Havugimana-Emili-2012 0.13%

A census of human soluble protein complexes. Havugimana et al (2012). *Cell*

Physical Interactions with 13,651 interactions from BioGRID

##### Huttlin-Harper-2017 0.11%

Architecture of the human interactome defines protein communities and disease networks. Huttlin et al (2017). *Nature*

Physical Interactions with 55,868 interactions from BioGRID

##### Brajenovic-Drewes-2004 0.06%

Comprehensive proteomic analysis of human Par protein complexes reveals an interconnected protein network. Brajenovic et al (2004). *J Biol Chem*

Physical Interactions with 118 interactions from BioGRID

##### Wang-Yang-2011 0.04%

Toward an understanding of the protein interaction network of the human liver. Wang et al (2011). *Mol Syst Biol*

Physical Interactions with 3,408 interactions from BioGRID

##### Huttlin-Gygi-2015 0.00%

The BioPlex Network: A Systematic Exploration of the Human Interactome. Huttlin et al (2015). *Cell*

Physical Interactions with 23,384 interactions from BioGRID

#### Co-expression 35.79%

##### Arijs-Rutgeerts-2009 4.23%

Mucosal gene expression of antimicrobial peptides in inflammatory bowel disease before and after first infliximab treatment. Arijs et al (2009). *PLoS One*

Co-expression with 676,695 interactions from GEO

#### Co-expression 35.79%

##### Bild-Nevins-2006 B 3.74%

Oncogenic pathway signatures in human cancers as a guide to targeted therapies. Bild et al (2006). *Nature*

Co-expression with 285,368 interactions from GEO

##### Alizadeh-Staudt-2000 3.08%

Distinct types of diffuse large B-cell lymphoma identified by gene expression profiling. Alizadeh et al (2000). *Nature*

Co-expression with 92,360 interactions from supplementary material

##### Burington-Shaughnessy-2008 2.97%

Tumor cell gene expression changes following short-term in vivo exposure to single agent chemotherapeutics are related to survival in multiple myeloma. Burington et al (2008). *Clin Cancer Res*

Co-expression with 295,320 interactions from GEO

##### Boldrick-Relman-2002 2.80%

Stereotyped and specific gene expression programs in human innate immune responses to bacteria. Boldrick et al (2002). *Proc Natl Acad Sci U S A*

Co-expression with 116,197 interactions from supplementary material

##### Rosenwald-Staudt-2001 2.78%

Relation of gene expression phenotype to immunoglobulin mutation genotype in B cell chronic lymphocytic leukemia. Rosenwald et al (2001). *J Exp Med*

Co-expression with 118,097 interactions from supplementary material

##### Roth-Zlotnik-2006 2.62%

Gene expression analyses reveal molecular relationships among 20 regions of the human CNS. Roth et al (2006). *Neurogenetics*

Co-expression with 683,844 interactions from GEO

##### Innocenti-Brown-2011 2.18%

Identification, replication, and functional fine-mapping of expression quantitative trait loci in primary human liver tissue. Innocenti et al (2011). *PLoS Genet*

Co-expression with 620,205 interactions from GEO

##### Perou-Botstein-2000 2.04%

Molecular portraits of human breast tumours. Perou et al (2000). *Nature*

Co-expression with 189,373 interactions from supplementary material

##### Dobbin-Giordano-2005 1.86%

Interlaboratory comparability study of cancer gene expression analysis using oligonucleotide microarrays. Dobbin et al (2005). *Clin Cancer Res*

Co-expression with 452,322 interactions from GEO

##### Chen-Brown-2002 1.70%

Gene expression patterns in human liver cancers. Chen et al (2002). *Mol Biol Cell*

Co-expression with 291,300 interactions from supplementary material

##### Wang-Maris-2006 1.64%

Integrative genomics identifies distinct molecular classes of neuroblastoma and shows that multiple genes are targeted by regional alterations in DNA copy number. Wang et al (2006). *Cancer Res*

Co-expression with 270,388 interactions from GEO

##### Wang-Cheung-2015 1.50%

Genetic variation in insulin-induced kinase signaling. Wang et al (2015). *Mol Syst Biol*

#### Co-expression 35.79%

##### Wang-Cheung-2015

Co-expression with 422,896 interactions from GEO

##### Ross-Perou-2001 0.92%

A comparison of gene expression signatures from breast tumors and breast tissue derived cell lines. Ross et al (2001). *Dis Markers*

Co-expression with 146,858 interactions from supplementary material

##### Jiang-de Kok-2017 0.85%

Omics-based identification of the combined effects of idiosyncratic drugs and inflammatory cytokines on the development of drug-induced liver injury. Jiang et al (2017). *Toxicol Appl Pharmacol*

Co-expression with 444,959 interactions from GEO

##### Ramaswamy-Golub-2001 0.60%

Multiclass cancer diagnosis using tumor gene expression signatures. Ramaswamy et al (2001). *Proc Natl Acad Sci U S A*

Co-expression with 284,829 interactions from supplementary material

##### Rieger-Chu-2004 0.25%

Toxicity from radiation therapy associated with abnormal transcriptional responses to DNA damage. Rieger et al (2004). *Proc Natl Acad Sci U S A*

Co-expression with 266,879 interactions from GEO

##### Perou-Botstein-1999 0.03%

Distinctive gene expression patterns in human mammary epithelial cells and breast cancers. Perou et al (1999). *Proc Natl Acad Sci U S A*

Co-expression with 68,200 interactions from supplementary material

#### Genetic Interactions 5.57%

##### IREF-SMALL-SCALE-STUDIES 3.19%

Genetic Interactions with 1,159 interactions from iRefIndex

##### Willingham-Muchowski-2003 2.01%

Yeast genes that enhance the toxicity of a mutant huntingtin fragment or alpha-synuclein. Willingham et al (2003). *Science*

Genetic Interactions with 37 interactions from BioGRID

##### Lin-Smith-2010 0.37%

A genome-wide map of human genetic interactions inferred from radiation hybrid genotypes. Lin et al (2010). *Genome Res*

Genetic Interactions with 4,805,334 interactions from supplementary material

#### Co-localization 2.92%

##### Johnson-Shoemaker-2003 2.51%

Genome-wide survey of human alternative pre-mRNA splicing with exon junction microarrays. Johnson et al (2003). *Science*

Co-localization with 426,464 interactions from GEO

##### Schadt-Shoemaker-2004 0.42%

A comprehensive transcript index of the human genome generated using microarrays and computational approaches. Schadt et al (2004). *Genome Biol*

Co-localization with 59,920 interactions from GEO

#### Shared protein domains 0.70%

##### INTERPRO 0.70%

#### Shared protein domains 0.70%

##### INTERPRO

Shared protein domains with 621,159 interactions from InterPro

#### Predicted 0.63%

##### Wu-Stein-2010 0.38%

A human functional protein interaction network and its application to cancer data analysis. Wu et al (2010). *Genome Biol*

Predicted with 89,967 interactions from supplementary material

##### I2D-INNATEDB-Mouse2Human 0.16%

InnateDB: facilitating systems-level analyses of the mammalian innate immune response. Lynn et al (2008). *Mol Syst Biol*

Predicted with 4,049 interactions from I2D

##### Stuart-Kim-2003 0.09%

A gene-coexpression network for global discovery of conserved genetic modules. Stuart et al (2003). *Science*

Predicted with 25,001 interactions from supplementary material

#### Pathway 0.55%

##### REACTOME 0.35%

Pathway with 24,890 interactions from Pathway Commons

##### NCI_NATURE 0.20%

Pathway with 10,118 interactions from Pathway Commons
